# StableMate: a statistical method to select stable predictors in omics data

**DOI:** 10.1101/2023.09.26.559658

**Authors:** Yidi Deng, Jiadong Mao, Jarny Choi, Kim-Anh Lê Cao

**Author notes:** indicates equal contribution (Jarny Choi, Kim-Anh Lê Cao).

## Abstract

Identifying statistical associations between biological variables is crucial to understand molecular mechanisms. Most association studies are based on correlation or linear regression analyses, but the identified associations often lack reproducibility and interpretability due to the complexity and variability of omics datasets, making it difficult to translate associations into meaningful biological hypotheses.

We developed StableMate, a regression framework to address these challenges through a process of variable selection across heterogenous datasets. Given datasets from different environments, such as experimental batches, StableMate selects environment-agnostic (stable) and environment-specific predictors in predicting the response of interest. Stable predictors represent robust functional dependencies with the response, and can be used to build regression models that make generalizable prediction in unseen environments.

We applied StableMate to 1) RNA-seq data of breast cancer to discover genes that consistently predict estrogen receptor expression across disease status, 2) metagenomics data to identify microbial signatures that show persistent association with colon cancer across study cohorts and 3) scRNA-seq data of glioblastoma to discern signature genes associated with development of pro-tumour microglia regardless of cell location.

Our case studies demonstrate that StableMate is adaptable to regression and classification analyses and achieves comprehensive characterisation of biological systems for different omics data types.

## 1 Introduction

Inferring relationships between biological variables is a critical problem in systems biology. Among different types of biological relationships, causal relationships are of high interest as they enable a deeper understanding of the function and regulatory mechanism of fundamental biological processes. However, this type of relationship is extremely difficult to identify based on observational studies alone without further investment in experimental design. In contrast, statistical associations (e.g. based on correlation or linear regression analyses) can be easily computed, but these associations may lead to spurious findings. In recent years, a large number of methods have been proposed for statistical association analysis. Most of theses methods are network modelling approaches that infer gene regulation by identifying associations between genes through their expression levels (Aibar et al., 2017; Chan et al., 2017; Chickering, 2002; Faith et al., 2007; Huynh-Thu et al., 2010; Langfelder and Horvath, 2008; Moerman et al., 2019; Spirtes et al., 2000). However, these approaches result in associations that are not robust against small variations in the data, are not reproducible or lack interpretability (Kang et al., 2021; Nguyen et al., 2021; Pratapa et al., 2020).

The concept of stability has gained popularity in recent years to improve the reproducibility and interpretability of conventional association analysis (Bühlmann, 2020). A statistical association is considered stable if it is invariant under small perturbations of the data, and hence is more likely to be reproducible across different studies or conditions. Stability analysis in this context allows us to gain unique biological insights that are not accessible with conventional inference association methods . Indeed, biological variables that show stable associations are more likely to be closely or even causally related in function compared to those with unstable associations. While stability is itself not sufficient to establish causality, a causal relationship is necessarily stable in some sense (Pearl et al., 2000). Thus, identifying stable associations may serve as a first step towards the inference on causal relationships. It is also enlightening to identify associations that are unstable as they are sensitive to a change of study and experimental conditions and provide insights into how these conditions influence a biological system (Shojaie, 2021).

We developed StableMate, an statistical framework to identify both stable and unstable associations through variable selection in a regression context. Inherent to variable selection is the motivation to infer regression function that encapsulates potential functional dependencies between the response and selected predictors beyond using simple correlations. StableMate is based on the recent theoretical development of stabilised regression (Pfister et al., 2021). Stabilised regression considers data collected from different ‘environments’ or experiments, including technical or biological conditions. Typical environments can be batches, cohorts, and also disease states. Given a response variable and a set of predictors measured on samples in multiple environments, there are two goals in stabilised regression. The first goal is to distinguish stable predictors from unstable predictors, based on whether these predictors are able to make consistent or inconsistent prediction of the response across multiple environments. The second goal is to build regression models using stable predictors that are generalisable to unseen environments. While the original approach from Pfister et al. (2021) provides an elegant framework for stabilised regression, its application is computationally inefficient for high-dimensional biological data. We showed in our simulation study that it can lead to inaccurate results. StableMate provides a new version of stabilised regression. While stabilised regression selects stable predictors by performing stability tests on every possible predictor subsets, StableMate optimises efficiency with a greedy search based on our improved stochastic stepwise selection algorithm.

We illustrate the broad applicability and flexibility of StableMate through three case studies across a broad range of biological questions and data types. We show that StableMate is able to 1) identify genes and gene modules involved in the trancriptional regulation of a critical breast cancer gene (Section 2.2), 2) identify fecal microbial markers for prediction of colon cancer while accounting for batch effects in a multi-cohort data (Section 2.3), and 3) characterise changes of microglia transcriptional identity during their transitions to a pro-tumour phenotype (Section 2.4). In both simulated and real data (Section 5, Supplementary Figure S2,S1), we benchmarked the prediction and the variable selection performances of StableMate against other commonly used regression methods, including the original stabilised regression algorithm in Pfister et al. (2021). The results show that StableMate yields superior performances compared to competing methods.

## 2. Results

### 2.1 Illustration of StableMate on toy example

We simulated 900 samples measured on 21 variables from three environments *e* = *e*^1^, *e*^2^, *e*^3^, with 300 samples in each environment. Denote by *Y* ^*e*^ the response variable of interest we wish to predict. We use the remaining 20 variables 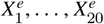 as predictors (see Figure 1A). In particular, 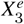 is a causal parent of *Y* ^*e*^ and is expected to be stable, whereas 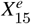 is a causal child of *Y* ^*e*^ and hence is expected to be environment-specific (see details in Supplementary Methods Section 7.2.1). To illustrate StableMate results we plotted *Y* ^*e*^ against 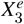 and 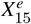 in Figure 1B. Here, 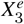 is stable in the sense that its relationship with *Y* ^*e*^ can be described by the same linear equation for *e* = *e*^1^, *e*^2^, *e*^3^, whereas *X*^15^ is environment-specific as its relationship with *Y* ^*e*^ varies for different *e*.

**Figure 1.**
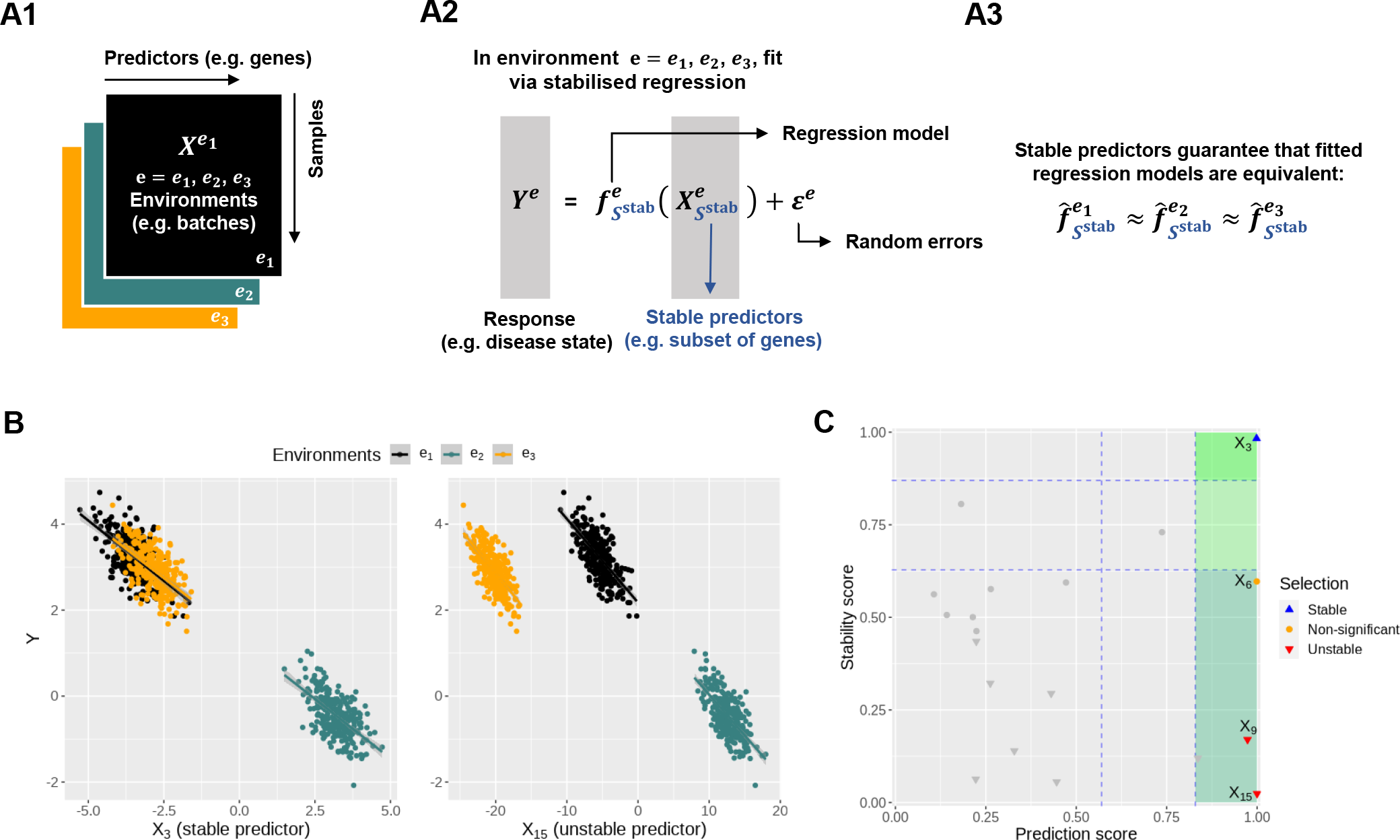
Toy example for StableMate analysis. **(A)** Stable predictors. Consider a regression problem where the response *Y* ^*e*^ and predictors *X*^*e*^ were generated from three different environments (e.g. batches, cohorts) *e* = *e*^1^, *e*^2^, *e*^3^, as represented in (A1). Stable predictors are a subset of all predictors that are useful for predicting *Y* ^*e*^ and whose association with the response *Y* ^*e*^ does not change with *e*. If we fit a regression model in each environment to predict the response using only the stable predictors (A2), then the fitted models should be approximately the same across all environments (A3). Thus identifying stable predictors is useful for constructing regression models that are agnostic to environment and hence may be more generalisable to unseen environments. On the other hand, predictive but unstable (referred to as’environment-specific’) predictors may be useful for understanding environment-specific regulatory mechanisms of the response *Y* ^*e*^. **(B)** Difference between stable and environment-specific predictors. We simulated 900 samples, each with response *Y* ^*e*^ and predictors 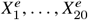 across environments *e* = *e*^1^, *e*^2^, *e*^3^. Left panel plots *Y* ^*e*^ against a stable predictor 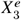; right panel plots *Y* ^*e*^ against an environment-specific predictor 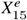 . Linear regression lines were fitted per environment. Both 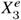 and 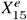 are useful for predicting *Y*^*e*^ since they are both strongly negatively correlated with *Y* ^*e*^ in each environment. However, for the stable predictor 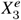, the regression lines have the same slope and intercept in all three environments. For the environment-specific predictor 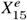, the regression lines have the same slope but differ in their intercepts. **(C)** StableMate variable selection plot. StableMate takes as input the predictors 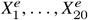 measured from the 900 samples across all environments, where the environment index *e* is known for each sample, and the response *Y* ^*e*^ for each sample. The variable selection plot shows the prediction score (x-axis) and the stability score (y-axis) assigned to each predictor. Vertical and the horizontal dashed lines represent the significance thresholds for prediction and stability respectively based on bootstrap of selections, as defined in the Method section. The predictive variables are further labeled as stable (blue) or environment-specific (red), where, in particular, 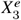 and 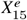 are both correctly labeled.

Briefly, the StableMate procedure first pools samples from all environments to identify the variables that are most predictive of the response regardless of the environment. Then, among these predictive variables, stable and unstable (environment-specific) variables are further differentiated. The variable selection results of StableMate can be summarised in Figure 1C, where every variable is assessed in terms of prediction and stability. First, all variables with low prediction scores (in gray) are filtered out. Second, among the predictive variables, we further differentiate between those that are significantly stable, unstable or indeterminate. In this example 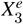 that expected to be stable received the highest prediction and stability score whereas 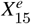 that is expected to be environment-specific received a high prediction score but the lowest stability score. We further detail in the Method Section 4.2 how we defined these scores and significance thresholds.

We evaluated the ability of StableMate to accurately identify stable and environment-specific predictors in a benchmark study where we compared our performance with existing approaches including the original stabilised regression algorithm. Our results show that StableMate leads to superior accuracy and computational efficiency, as detailed in Supplementary Figures S2,S1.

The next sections highlight the flexibility of StableMate to identify stable and environment-specific predictors in different analytical settings. The different types of analyses are described in Table 1.

**Table 1.**
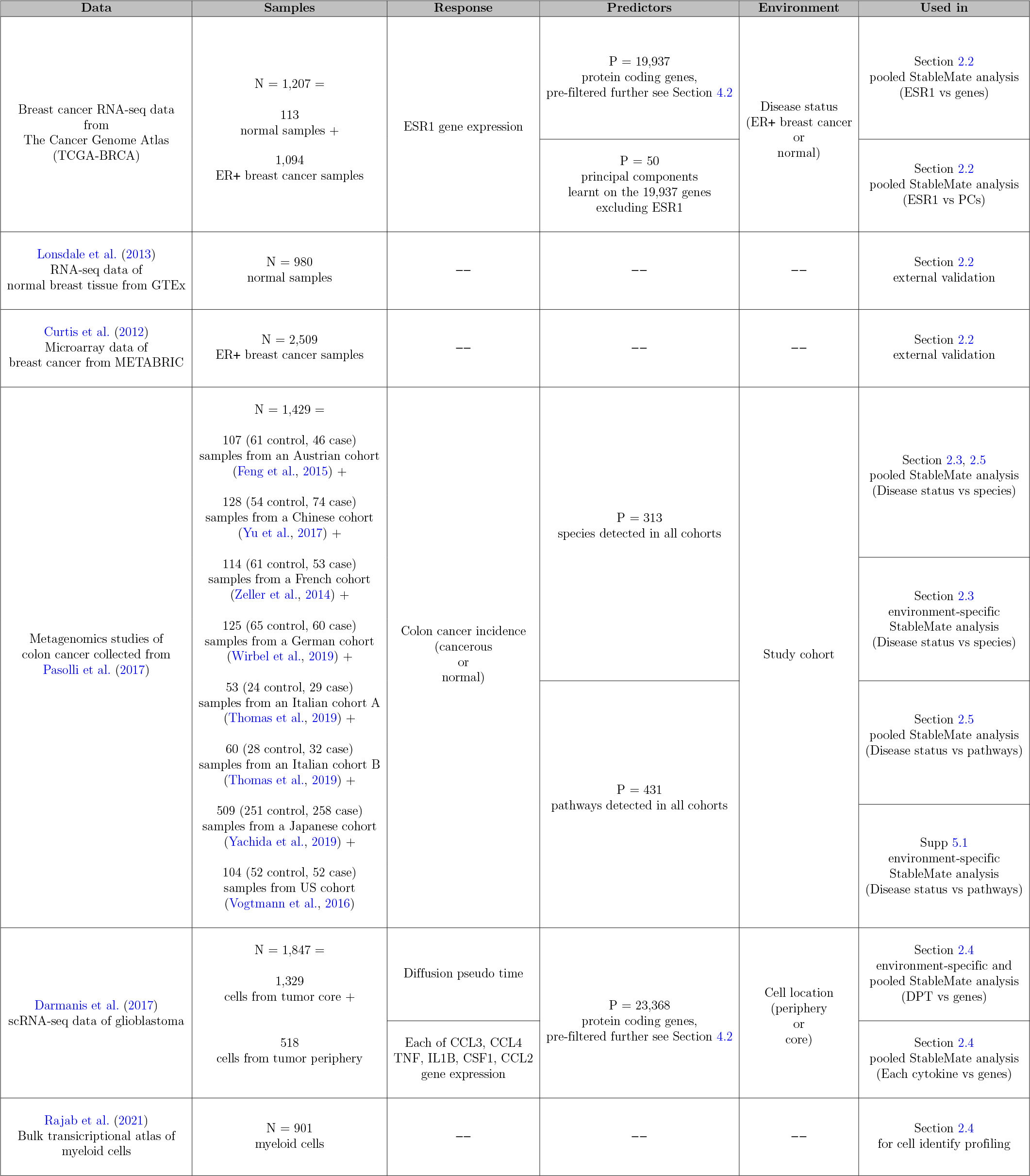
Summary of case studies. Sample break down per environment, response, predictors and the environment variables are described for StableMate regression are described. We performed two types of StableMate analysis based on how predictive variables were defined. In the first type, we pooled environments to select predictive variables and assess their stability across environments. In the second type, we select predictive variables in each environment and tested the stability of the predictor selected in the remaining combined environments. These two types of StableMate analysis are referred to as *pooled* and *environment-specific*.

### 2.2 StableMate identifies genes associated with ESR1 expression in ER+ breast cancer using RNA-seq data

In this first case study, we used StableMate to identify genes and gene modules associated with regulation of the ESR1 gene based on transcriptomic data. ESR1 is one of the marker genes of the ER+ subtype of breast cancer (BC), characterised by the high expression of the estrogen receptor (ER) (Johnson et al., 2021). We are interested in the association between ESR1 and other genes across normal and ER+ samples. In particular, we expect that genes identified as stable for predicting ESR1 expression are not confounded by disease status, suggesting close or potential causal relationship with ESR1 in its transcriptional regulation. In contrast, genes identified as disease-specific might be interacting with ESR1 indirectly, e.g, at the downstream of ER regulation or by co-regulating with ER.

#### Data and StableMate setting

We used the publicly available BC gene expression (BRCA) data from The Cancer Genome Atlas (TCGA, Weinstein et al. (2013)). We filtered the dataset to retain 113 normal and 778 ER+ tumour samples.

Since we were interested in the regulation of ESR1, we set ESR1 as the response and all other genes as predictors. We set the disease status (normal or ER+) as the environment variable, so that we could identify stable genes, whose association with ESR1 did not change significantly between normal vs. ER+ samples, and disease-specific genes, whose association with ESR1 significantly varied significantly between disease status. In addition to identifying individual genes, we also combined StableMate with principal component analysis (PCA) to identify stable and disease-specific gene modules. Namely, we still took ESR1 as the response and disease status as the environment variable, but we used metagenes (first a few PCs of all genes except ESR1) as predictors. Then, we defined the stable and disease-specific gene modules as the the most important genes of stable and disease-specific meta-genes.

#### StableMate selected genes proxy to ESR1 regulation

The StableMate variable selection results are summarised in Figure 2A. Among the most stable genes predictive of the ESR1 expression, CCDC170 and ARMT1 are the closest genes located to the upstream genomic region of ESR1 (Supplementary Figure S3A) and have been reported to fuse with ESR1 (Vitale et al., 2022). Their proxy to ESR1 suggests that they might be subject to the same transcriptional regulation as ESR1, thus explaining their stability. On the other hand, the STC2 gene was identified as disease-specific. This might be explained by the fact that STC2 has been identified as a downstream target of the ER signaling (Bouras et al., 2002; Raulic et al., 2008). In addition, the proximal promotor region of STC2 is not directly subject to ER binding but is dominated by other transcriptional activities such as hypoxia induced stress response (Law et al., 2008; Law and Wong, 2010). As a result, ER signaling is indirectly involved in the STC2 activation (Raulic et al., 2008). This evidence supports our hypothesis that STC2 and ESR1 should be indirectly or distally related in transcriptional regulation as indicated by their environment-specific associations.

**Figure 2.**
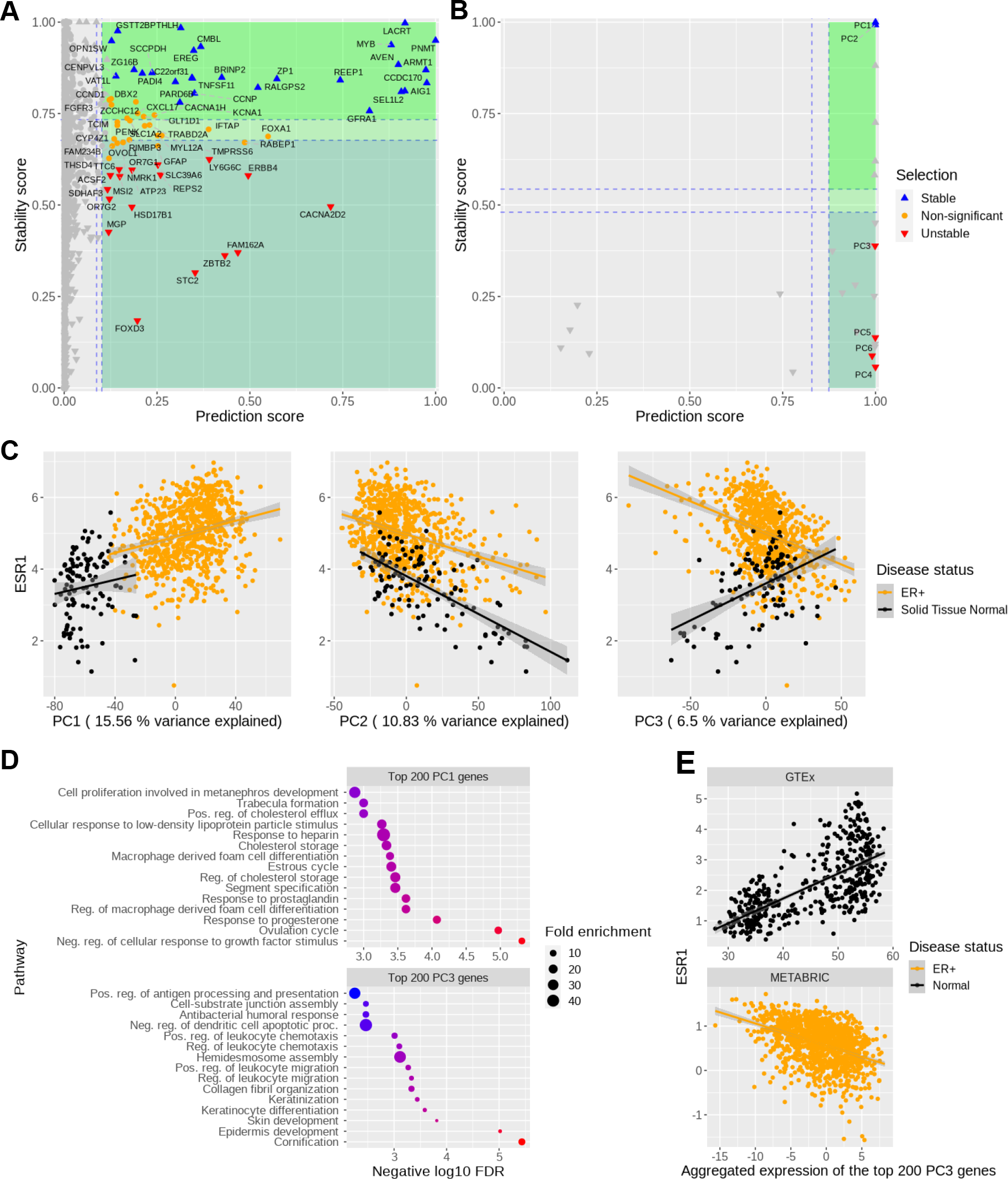
StableMate selects genes from TCGA-BRCA dataset which predict ESR1 expression across normal and ER+ samples. We used **(A)** gene expression or **(B)** principal components (PCs) of gene expression as predictors. The stability score (y-axes) of a gene is a measure of how consistently this gene predicts ESR1 regardless of the disease status (normal or ER+). Stable and disease-specific genes/PCs are colored in blue and red, respectively. **(C)** ESR1 expression (y-axis) against PC scores (x-axis). The correlation between ESR1 with the highly disease-specific PC3 changed from positive to negative between normal and ER+ samples, whereas the sign of the correlations between ESR1 and the stable PC1 and PC2 remained unchanged between normal and ER+ samples. We analysed PC1 (i.e. the most important stable PC) and PC3 (i.e. the most important disease-specific PC) as an example. **(D)** Gene ontology enrichment on the top 200 genes from PC1 (top) and PC3 (bottom) suggested biological activities related to hormonal regulation and epidermis development, respectively. The predictive ability and stability of PC1 suggest that ESR1 may directly participate in hormonal regulation, which is corroborated by the knowledge that ESR1 is a transcriptional factor activated by the estrogen binding. **(E)** Reproduciblility of StableMate results using external databases, GTEx for normal breast tissue and the METABRIC data from cBioPortal for ER+ BC: ESR1 expression against the expression of the metagene defined by the top 200 genes contributing to PC3 (i.e, linear combination of these 200 genes according to the loading vector of PC3) confirm the opposite trends we observed in **(C)** of ESR1 against PC3 in normal and ER+ samples.

#### StableMate with PCA identified gene modules associated with ESR1

Feature selection from transcriptional data is often followed by gene set enrichment analysis. While the stability analysis on individual genes gave us some insights into the ER regulation here, it selected relatively few genes as either stable or disease-specific - insufficient for statistically meaningful enrichment analysis. To overcome this issue, we used the first 50 PCs of all genes (except ESR1) as predictors for ESR1 expression rather than individual genes. In this context, each PC is a linear combination of the expression of all genes except ESR1, and can be viewed as a meta-gene, which is useful for quantifying the activities of gene modules (Langfelder and Horvath, 2007). Similar to our previous analysis, disease status (normal and ER+) was set as the environmental variable.

The StableMate variable selection results are summarised in Figure 2B. PC1 and PC2 were found to be highly stable and predictive, suggesting that the major source of variation they explain (15.56% and 10.83% respectively) is closely related to the ER regulation. All subsequent PCs up to PC6 were predictive but disease-specific. We considered the top 200 genes contributing to PC1 (most stable) and PC3 (disease-specific) and conducted an enrichment analysis. Genes from PC1 were mainly associated to biological processes related to hormone regulation (Figure 2C). The ESR1 mediated estrogen signaling is at the center of hormone regulation, and hence the high prediction ability and stability of PC1 is manifest. Genes from PC3 were associated with basal cell like transcriptional activities in epidermis development. The top genes contributing to PC3 (see details in Supplementary Figure S3B), included a high proportion of basal cytokeratins (BCKs), such as KRT5, KRT7, KRT14 and KRT17, suggesting that PC3 may reflect the ‘basalness’ of samples (Figure 2C). Interestingly, PC3 scores were positively correlated with ESR1 expression in normal samples but negatively correlated with ESR1 in ER+ samples (Figure 2C). This trend was also observed between the basal BC enriched genes (listed in Li et al. (2022)) and ESR1 expression (Supplementary Figure S3C), confirming the PC3 characterisation of basalness.

To validate the reproducibility of our findings, we queried the gene expression portals GTEx (Lonsdale et al., 2013) for normal breast tissue and the METABRIC data from cBioPortal (Cerami et al., 2012) for ER+ BC. Our analysis using these external datasets showed similar trends between ESR1 and PC3 (Figure 2E). The negative correlation between BCKs (contributing to PC3) and ESR1 expression may be explained by the fact that the BCK induction in ER+ BC requires low ER expression (Li et al., 2022). However, to the best of our knowledge, no study so far has reported that this correlation may turn positive in normal breast tissues.

### 2.3 StableMate discerns global microbial signatures for colon cancer in multi-cohort metagenomics data

There has been considerable research interest in using fecal microbiome as biomarkers for colorectal cancer (CRC). If successful, this non-invasive way of screening for CRC may reduce the mortality rate through early intervention (Labianca et al., 2010; Sears and Garrett, 2014). By pooling fecal metagenomics data from a large number of independent CRC–control studies, several meta-analyses have been conducted to identify cross-cohort microbial signatures of CRC and to build predictive models for its diagnosis (Dai et al., 2018; Thomas et al., 2019; Wirbel et al., 2019). However, these analyses ignored the technical differences between cohorts, which could have confound their results. We addressed this problem by conducting a meta-analysis based on StableMate using the cohort as the environmental variable. In particular, we selected stable microbial signatures that make consistent predictions of CRC across cohorts, as well as cohort-specific signatures that highlight confounding factors in CRC prediction. Our results showed better prediction accuracy compared to the methods used in these studies.

#### Data and StableMate setting

We retrieved eight CRC case-control fecal metagenomic datasets from the R package *curatedMetagenomicData* (Pasolli et al., 2017). The datasets were generated by eight different cohorts from seven countries (refer to Table 1 for the cohort used and for the number of CRC and controls in each cohort). Data were curated into abundance data using a standardised data processing pipeline by Pasolli et al. (2017). In total, we collected 604 CRC and 596 control samples. Our analysis focused on the species abundance data measured on 313 microbial species. The analysis of pathway abundance data measured on 431 pathways is detailed in Supplemental Figures S4 and S6.

Since we were interested in the CRC diagnosis using metagenomics data, we set the disease status (CRC or normal) as the response and the microbial species as predictors. We implemented StableMate using the following two strategies. In a first analysis, we applied StableMate as in the toy example (Section 2.1) and our first case study (Section 2.2), where all cohorts were pooled to select predictive species and assessed their stability by setting cohort as the environment variable. From this analysis, we found that the majority of the selected predictive species were stable and none of them was cohort-specific. Therefore, in a second analysis, we applied StableMate on each cohort to identify cohort-specific predictive species, and tested the stability of the species selected in the remaining seven cohorts combined. There we considered only two environments: the specific cohort and the remaining cohorts combined. These ‘cohort-specific analyses’ are useful for identifying species that are highly predictive in a specific cohort but their association with CRC in the specific cohort cannot be generalised to other cohorts.

#### StableMate identified stable microbial species predictive of colon cancer

From the pooled analysis, we identified 23 stable species to predict disease status (CRC or normal) (Figure 3A). To assess these stable species, we compared the principal coordinate analysis (PCoA) using all 313 species versus the 23 stable species. The PCoA results combined with a permutation ANOVA showed that the main source of variation was the cohort effect rather than disease status (Figure 3B1) when all species were used. This implies that a predictive model built using all species is likely to be affected by cohort (batch) effects. In contrast, the PCoA and ANOVA results of the 23 stable species selected by StableMate showed a decrease in cohort effects and an increase in the effects of disease status (Figure 3B2). In particular, the CRC and normal samples were better separated in the PCoA when using only the 23 stable species (left panel of Figure 3B2).

**Figure 3.**
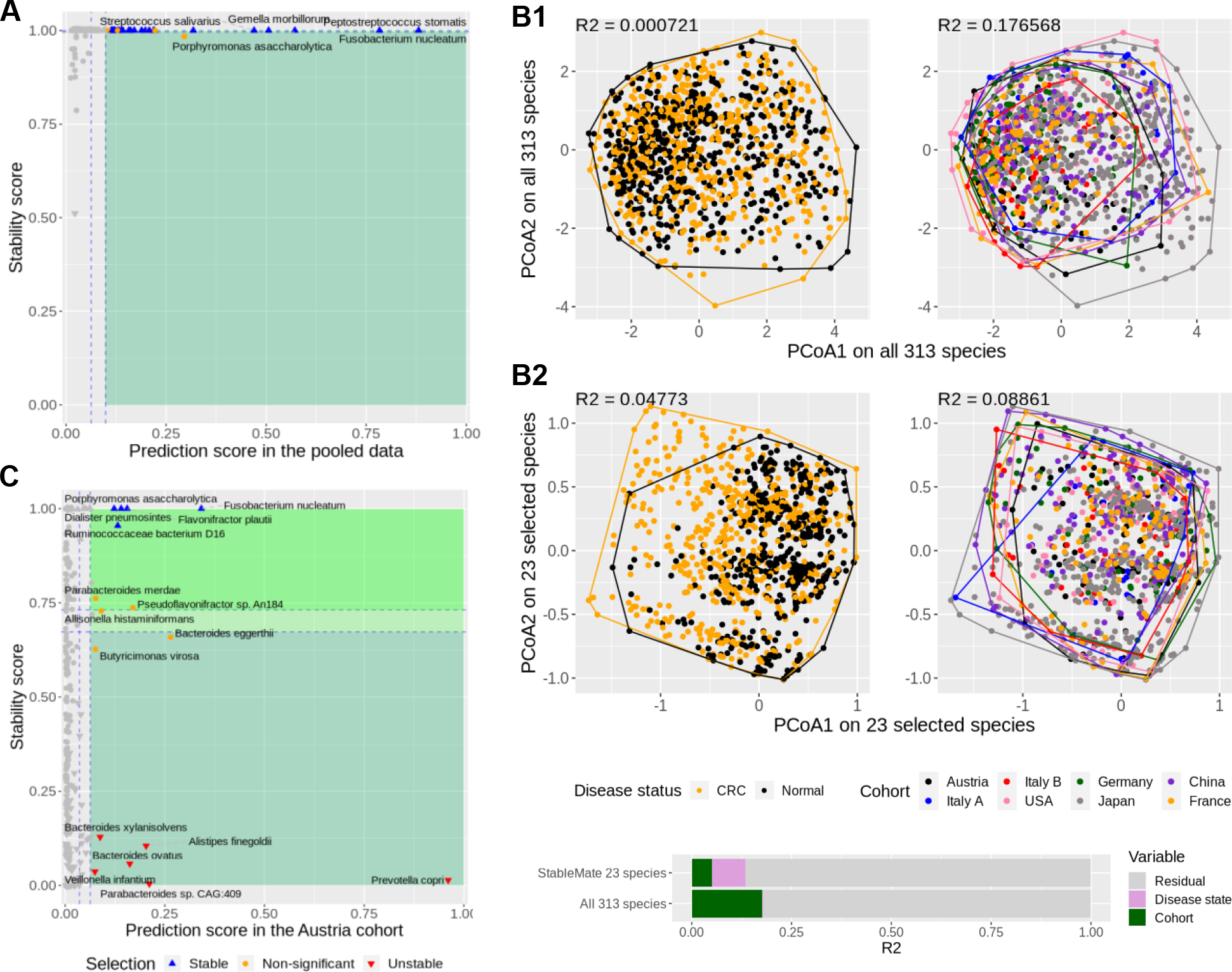
StableMate meta-analysis of metagenomic data reveals key species predictive of CRC across eight independent study cohorts. **(A)** StableMate variable selection plot of the pooled analysis. The majority of highly predictive species were found stable and none was identified as cohort-specific. **(B)** Principal coordinate analysis (PCoA) with samples coloured by either disease status (left column) or cohorts (right column). **B1**: using all 313 species shared by all cohorts, regardless of their stability; **B2**: using only the 23 stable species selected by StableMate. Permutation ANOVA *R*^2^ statistics on the first two principal coordinates is shown on the top-left corner of each panel. The coloured bar at the bottom shows the composition of the total variance. When considering all 313 species, the cohort effect is much larger than the disease effect (almost negligible); with 23 species identified as stable, the cohort effect is still present but smaller than the disease effect. **(C)** StableMate variable selection plot of the Austria cohort-specific analysis (one of the eight cohort-specific analyses). *Prevotella copri* was found to be an Austria-specific species for predicting CRC, since it has a high prediction score but a low stability score. Such species are interesting for studying cohort-specific effects that may confound the CRC diagnosis.

#### StableMate identified cohort-specific microbial species predictive of colon cancer

We conducted cohort-specific analyses for each of the eight cohorts to identify predictors with high cohort specificity. As an example, Figure 3C shows the results for the Austrian cohort. A number of species were found to be highly predictive in the Austrian cohort but with a very low stability score, and therefore were identified as cohort-specific. Among them, *Prevotella copri* was the most predictive and one of the most cohort-specific species, suggesting that *Prevotella copri* might be a marker for CRC specific to the Austrian cohort only. It should be noted however, that diet could be a confounder, as the Austria-specific species might be related to the low-fiber diet in that population (see Supplementary Results 5.1 for details).

#### Pooling data improves generalisability of prediction models

Most of the predictive species selected on the pooled data were stable (Figure 3A), whereas predictive species selected in the individual cohorts showed less stability (Supplementary Figure S5). This is expected since training regression models on pooled data can yield improved generalisability compared to training on individual datasets as shown in the other meta-analysis studies (Dai et al., 2018; Thomas et al., 2019; Wirbel et al., 2019). However, aside from pooling data, we were able to further improve generalisability of prediction models by taking into account the cohort effect through stability analysis. We conducted a benchmark study in Section 2.5, where we showed that the StableMate model built using the stable species outperformed several commonly used regression methods in the pooled analysis.

### 2.4 StableMate characterises cell identity transition of glioblastoma associated microglia with scRNA-seq data

Glioblastoma (GBM) is the most invasive type of brain tumour that presents significant therapeutic challenges. GBM harbors a heterogeneous tumour microenvironment dominated by Tumour-Associated Macrophages (TAM) and microglia, which were recruited by GBM to promote tumour growth, migration, recurrence and resistance to immunotherapy (Andersen et al., 2021). Since the majority of TAM in GBM are thought to be derived from microglia (i.e. tissue-resident macrophages in the brain) infiltrating the tumour, identifying key genes involved in this process could have therapeutic potential.

In this case study, we analysed a single cell RNA sequencing (scRNA-seq) dataset of myeloid cells at the periphery (migrating front) and the core of the GBM tumour. These locations represent the start and the end points of the transition from microglia to TAM. We used StableMate to extract the key genes involved in this transition, while taking location into account. Hence we were able to investigate how does the transition differs between the locations and reveal location-agnostic and -specific immune activities

#### Data and StableMate setting

From the scRNA-seq dataset from Darmanis et al. (2017) of four GBM tumours, we extracted and analysed 1,847 myeloid cells from the tumor core (1,329 cells) and from the tumor periphery (518 cells) with the cell annotation provided by the authors of the study.

We visualised the scRNA-seq data and observed a clear cell trajectory between the two locations, which may represent the celluar transition of microglia to TAM. We conducted a pseudo-time analysis to quantify this trajectory, and used StableMate to predict as a response the pseudo-time based on expression of the genes as predictors. The cell location, core or periphery, was set as the environment variable. StableMate selected several cytokines as being predictive of the pseudo-time. To further investigate the possible mechanism of these cytokines, we performed a second analysis to build a gene regulatory network for these cytokines. More specifically, we applied StableMate on each of the cytokines as response and all other genes as predictors. We then summarised the selection results in the form of a network, where the cytokines are connected to their stable predictor genes.

#### Diffusion pseudo-time tumour periphery to core

We first visualised the myeloid cells projected onto a bulk RNA-seq reference atlas of myeloid cells (Rajab et al., 2021) to assign the identity of the cells. We performed this using Sincast (Deng et al., 2022) (Figure 4A1). The projection showed that the cells from the periphery of the tumour closely matched fetal microglia in the reference, and the cells from the core of the tumour matched a wider range of monocytes and macrophages (Supplementary Figure S8A). The projection also showed a continuous state of transition, rather than discrete clusters. We also confirmed the transition by a separate diffusion map analysis in Supplementary Figure S8B. This exploration suggested that the data were suitable for diffusion pseudo-time (DPT) analysis (Figure 4A2), where we set the cells at the tumour periphery as the root (start) of the trajectory (Haghverdi et al., 2016). The inferred DPT was then used as response for our first StableMate analysis described below.

**Figure 4.**
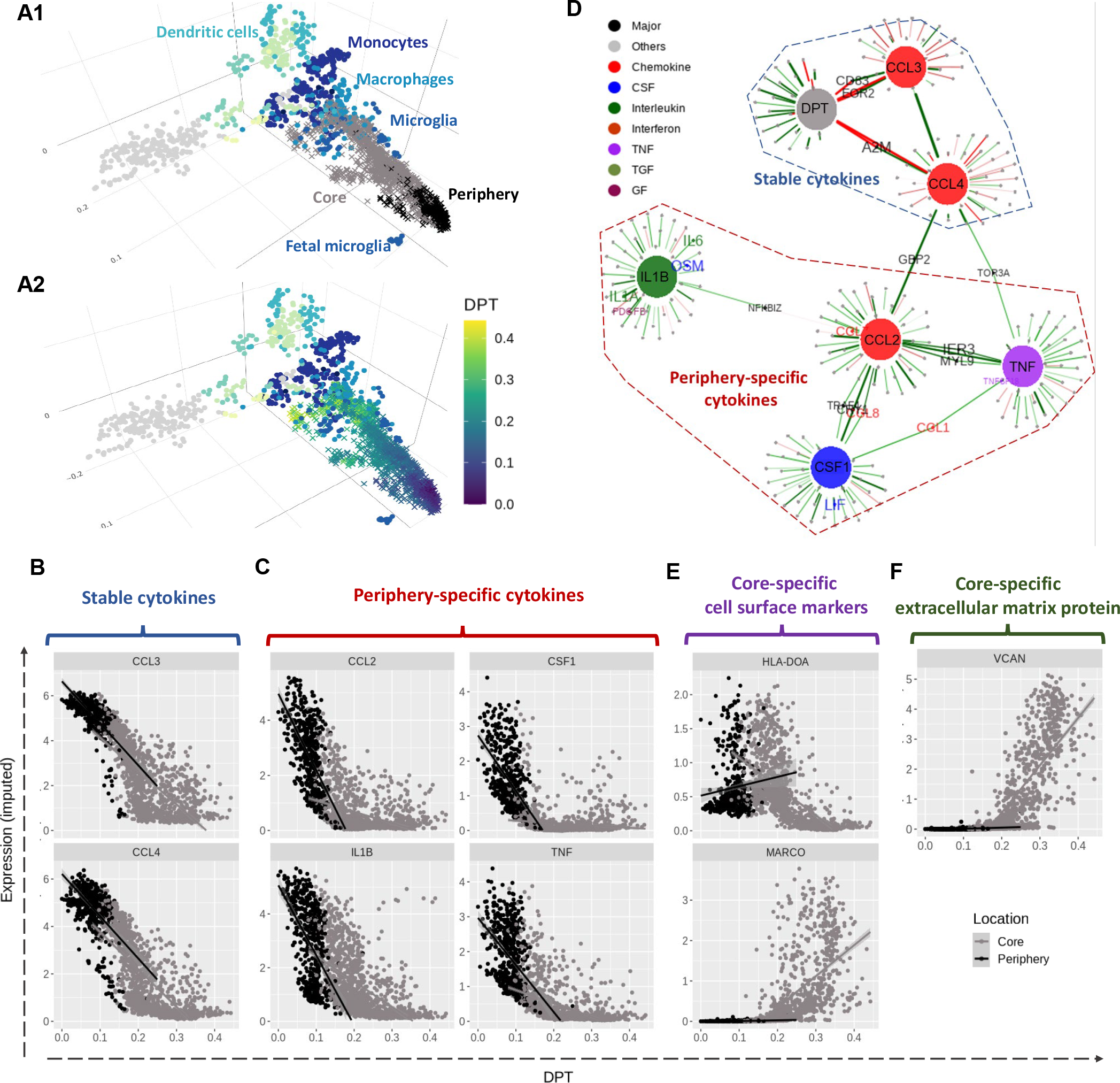
Characterising transition of microglia cell identity from periphery to core in glioblastoma tumour with scRNA-seq data. **(A1)** Sincast projection of the query single cells (crosses) onto a bulk RNA-seq reference atlas of myeloid cells (dots) to assign cell identity. The cells from the tumour periphery were located close to the reference fetal microglia, while the cells from the tumour core showed a transition towards the reference monocytes and macrophages. **(A2)** is identical to (A1) except that cells are coloured according to Diffusion Pseudo-Time (DPT), representing a cell state transition. StableMate was applied to select genes predictive of DPT, where cell location (core and periphery) was set as the environmental variable. **(B)-(F)** The expression of the cytokines was imputed based in Sincast. We identified several cytokines that are typical microglia activation and polarisation markers, including **(B)** CCL3 and CCL4, which are stable and **(C)** TNF, IL1B, CCL2 and CSF1, which are periphery-specific. **(D)** A gene regulatory network was built by running StableMate on each of the seven response variables, namely DPT and six cytokines CCL3, CCL4, TNF, IL1B, CCL2 and CSF1 (represented as large nodes). The aim was to select stable and predictive genes associated with each of these response variables. The cell location was still set as the environment variable. An edge indicates that a gene is stable and predictive of a response variable. We found that CCL3 and CCL4 were stable and predictive of DPT as a separate graphical community than TNF, IL1B, CCL2 and CSF1, which were predictive but unstable of DPT. **(E)** The expression levels of MHC-II molecule HLA-DOA and the macrophage marker MARCO. **(F)** The expression levels of large extracellular matrix protein VCAN. MARCO, VCAN and HLA-DOA were all identified as core-specific. The up regulation of MARCO, VCAN and the down regulation of HLA-DOA suggests a development of M2-like immunosuppressive macrophage.

#### StableMate analysis identifies cytokines that signify microglia pre-activation and polarisation in tumor periphery

Among the genes selected by StableMate as predictive of DPT, we identified six cytokines whose expression were all negatively correlated with DPT (Supplementary Figure S8C). Amongst these cytokines, CCL3 and CCL4 were identified as stable (Figure 4B), while TNF, IL1B, CCL2 and CSF1 were identified as periphery-specific (Figure 4C). The selection of the six cytokines are interesting as they are important markers of microglia activation in response to disease (Jurga et al., 2020).

In order to visualise the relationships of these cytokines, we ran StableMate on each cytokine as a response, where all the other genes were used as predictors, and built a gene regulatory network (Figure 4D - DPT was included as a ‘pseudo gene’ here to incorporate the result from the first analysis). This network showed that the two stable cytokines (CCL3 and CCL4) formed a community with DPT, whereas the four periphery-specific cytokines (TNF, IL1B, CCL2 and CSF1) formed another community.

Other stable genes predictive of DPT represented on this network include EGR2 and CD83 which were connected to both DPT and CCL3 (Supplementary Figure S8D). CCL3, CCL4, CD83 and EGR2 are all known to be associated with immediate early inflammatory response by microglia in a pre-activated state, which are in between homeostasis to those fully activated under pathological conditions (Kohno et al., 2014; Masuda et al., 2019, 2020; O’Donovan et al., 1999; Sinner et al., 2023; Veremeyko et al., 2018). These four genes showed consistent down-regulation during the transition regardless of cell location. On the contrary, the periphery-specific cytokines, which are known markers for microglia polarisation to either the pro-inflammatory M1 or anti-inflammatory M2 phenotype (Jurga et al., 2020), exhibited stronger negative association with DPT in the periphery - resulting in low expression levels, and weak association with DPT in the core. (Figure 4C).

#### Core-specific genes revealed reprogramming of tumour-infiltrating microglia into immunosuppressive TAM in GBM tumours

The core-specific genes identified by StableMate included two interesting cell surface markers: the marcophage marker MARCO and MHC class II antigen HLA-DOA (Supplementary Figure S8C). MACRO was lowly expressed in the tumour periphery but up-regulated along the DPT trajectory towards the tumor core (Figure 4E upper). HLA-DOA expression levels had a low-high-low pattern along the trajectory, with high expression levels at the boundary of tumour periphery and core (Figure 4E lower, other MHC-II genes were also examined in Supplementary Figure S8E). The up-regulation of MARCO and the down-regulation of HLA-DOA towards the core may indicate the presence of MARCO^*hi*^, MHC-II^*lo*^ macrophages, which are characteristic of the M2-like immunosuppressive TAM (Georgoudaki et al., 2016; Wang et al., 2011). In addition to these two cell surface markers, many pro-tumour markers were also identified by StableMate as core-specific and showed similar expression patterns as MACRO (Supplementary Figure S9A). One example is VCAN, which encodes a large extracellular protein contributing to the establishment of tumour microenvironments (Figure 4F). The expression patterns of these core-specific pro-tumour markers suggest that they responded specifically to the tumour microenvironment and hence are potentially good therapeutic targets.

In addition, we examined the immune activation state of the cells at the beginning of the core stage of the transition. We observed high expression of the stable cytokines CCL3 and CCL4 (Figure 4C), as well as the microglia marker TMEM119, which were all then gradually suppressed in the core (Supplementary Figure S8F). This may imply the reprogramming of activated microglia in the early stages of the core-transition to TAMs.

### 2.5 Benchmarking StableMate variable selection and prediction on metagenomics data

We used the species abundance data from eight metagenomics studies of CRC described in Section 2.3 to benchmark the variable selection and prediction performances of StableMate (using logistic regression model as the base model) against Generalised Linear Model (GLM with logistic regression using all predictors), Lasso regression (Tibshirani, 1996) and random forest (RF, Breiman 2001). To assess the prediction performance of these methods, we used a leave-one-dataset-out (LODO) cross-validation strategy. That is, in each of the eight cross-validation iterations, we left out one of the cohort and trained the different regression models using the other seven cohorts (based on all 313 species). The left-out cohort was then used as a test dataset, on which the area under the receiver operating characteristics curve (AUC) was calculated for each regression model. Since the left-out cohort represents an unseen environment, regression models receiving higher AUC can be considered as more generalisable. To assess the variable selection performance of StableMate, recall that we have already applied StableMate to do a pooled meta-analysis as described in Section 2.3 and identified 23 stable species. We applied Lasso and RF to the same pooled data with eight cohorts to select 23 species (for RF, we ranked all species by their importance scores in descending order, and then selected the top 23). We use these three lists of species to build RF models and assess their generalisability using LODO.

#### StableMate outperformed commonly used regression models in classifying CRC

The LODO AUC values for all competing methods are shown in boxplots in Figure 5A. To illustrate the benefits of using stable predictors to build regression models, we considered two versions of the StableMate prediction model, one built using all selected predictive variables, the other using only the stable predictors. To further investigate if the differences in AUC values were statistically significant, we conducted a series of two-sided paired t-tests and the p-values of these tests are shown on Figure 5A.

**Figure 5.**
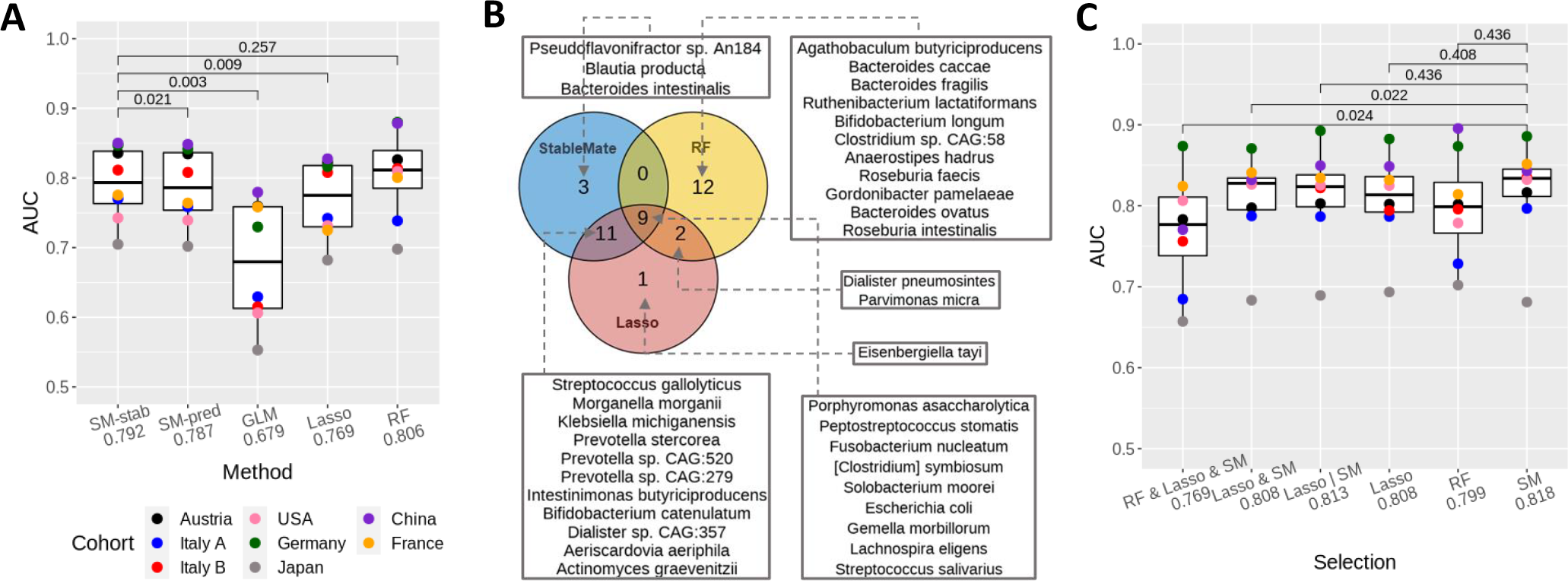
StableMate outperforms commonly used regression methods in prediction and variable selection based on the colon cancer case study. **(A)** We used leave-one-dataset-out (LODO) cross-validation to calculate area under the ROC curve (AUC, y-axis) and assess the generalisability of the classification when applied to an unseen cohort. Paired t-tests compare the AUC values and adjusted p-values (Benjamini and Hochberg, 1995) are shown . Each point presents the AUC value calculated on a left-out cohort. Methods include GLM: generalised linear model (logistic regression), Lasso: Lasso logistic regression, RF: random forest and two versions of StableMate (logistic regression), SM-Stab based stable predictors only and SM-Pred using all predictive variables. Among all linear methods (all except RF), SM-Stab obtained the highest mean AUC (difference is statistically significant). Compared to RF, SM-Stab had slightly lower mean AUC, but this difference was not statistically significant. Note that RF is a more flexible non-linear classification method. **(B)** Venn diagram to compare the three lists of species (each containing 23 species) selected by StableMate, Lasso and RF. StableMate and Lasso made similar selections, with 20 species selected by both. The RF selection was quite different to the other two methods. Nine species were selected by all three methods, all of which are known to be associated with CRC (Ternes et al., 2020). In addition, two species, also known to be associated with CRC, were selected by both Lasso and RF but not by StableMate. This is because these two species were not significantly stable as suggested by StableMate selection. **(C)** Generalisability of six sets of species: top 23 species selected by StableMate (‘SM’), Lasso and RF, the 9 species selected by all the methods (‘RF & Lasso & SM’), the 20 species selected by both Lasso and StableMate (‘Lasso & SM’) and the 26 species selected by either Lasso or StableMate (‘Lasso | SM’). We built six RF classifiers using these six sets of species and reported their AUC values (mean AUC on the x-axis). The stable species selected by StableMate led to the the best RF model, with a higher AUC than RF trained with all 313 species in (A).

We first compared the performances of all methods except RF, since they all use variants of linear models to make prediction, whereas RF is a nonlinear approach. From Figure 5A we observed that the StableMate prediction using only stable predictors was significantly better than GLM, Lasso and StableMate using all predictive variables. Among these methods, GLM had the worst generalisability. As GLM does not perform variable selection, the prediction model potentially included many noisy features. Lasso’s selected variables led to poorer prediction compared to the two variants of StableMate. The version of StableMate using only stable predictors led to slightly higher mean AUC compared to StableMate using all predictive variables, with a p-value indicating a significant difference. The superior performance of StableMate based on stable predictors highlighted the benefits of using such type of predictors to build prediction models.

Finally, we observed that the AUC performance of StableMate based on the stable predictors was indistinguishable from RF. This can be explained as StableMate only uses an ensemble of stringent logistic models to make predictions (see Method section 4.2), whereas RF uses an ensemble of nonlinear and highly flexible decision trees targeted for classification tasks. The notable classification performance of RF motivated our second benchmark study below, in which we assessed the generalisability of species selections by evaluating RF models built on these selections.

#### Species selected by StableMate lead to more generalisable prediction models

We applied StableMate, Lasso and RF to select three species lists, each containing 23 species. Figure 5B shows a Venn diagram that compares these three lists (see also Supplementary Figure S7A for a more comprehensive comparison). The three lists included an overlap of nine species, all of which are well-known species associated with CRC (Ternes et al., 2020). Among these three methods, StableMate and Lasso shared 20 species, many more than with RF species selection. For a quantitative comparison, we built RF models using six different selections of species and computed their AUC using the LODO cross-validation approach. The results are summarised in Figure 5C, where the 23 stable species selected by StableMate led to the highest mean AUC (0.818). StableMate and Lasso selections had high AUC values, since their selections were similar. The difference between StableMate and RF selections was not statistically significant, probably due to a lack of cohorts and statistical power. However, the StableMate selection led to less variable prediction performances (smaller interquartile range) compared to RF.

Of note, two species, *Dialister pneumosintes* and *Parvimonas micra*, were selected by Lasso and RF but not by StableMate (Figure 5B). In particular, *Parvimonas micra* is known to be associated with CRC as it promotes tumourigenesis (Chang et al., 2023; Zhao et al., 2022). However, StableMate identified these two species as predictive but not significantly stable. The fact that there was no improvement in prediction performance of RF trained on the StableMate selection with these two additional species (comparing ‘Lasso | SM’ and ‘SM’ in Figure 5C) justifies why StableMate did not select these two species.

A similar benchmarking analysis based on the pathway abundance data showed that StableMate outperformed the other methods, including RF, in predicting CRC (hightest mean AUC, see Supplementary Figure S6B). However, the pathway abundance data were less stable and less predictive of CRC compared to the species abundance data. All methods obtained lower AUC scores in LODO assessment. We observed strong differences in variable selections between the cohorts and the methods (Supplementary Figure S6A, S7).

## 3 Discussion

The unbiased characterisation of a biological system requires a comprehensive understanding of the relationships between biological variables. Current methods that infer biological relationships attempt to define and identify statistical associations but often lack generalisability or biological interpretability (Kang et al., 2021; Nguyen et al., 2021; Pratapa et al., 2020). We developed StableMate, a new regression framework based on stabilised regression (Pfister et al., 2021) to address these challenges.

StableMate selects stable and environment-specific (unstable) predictors of the response variable to represent statistical associations across different technical or biological environments. Discerning stability of associations allows us to make interpretable inference on biological relationships. On the one hand, stable predictors suggest closer relationships with the response compared to environment-specific predictors. On the other hand, environment-specific predictors are useful for characterising the environmental differences on the biological system under study.

In the three case studies dealing with different types of cancer omics data, we showed that StableMate brings novel biological insights. In the simulation study, we showed the benefit of using StableMate for better prediction accuracy and computational efficiency, and accuracy of variable selection compared to existing methods.

In the first case study, we analysed RNA-seq data of breast cancer. Stability analysis allowed us to identify genes and gene modules that directly or indirectly relate to ESR1 regulation.

In the second case study, we conducted a meta-analysis of eight metagenomic studies of colon cancer. StableMate analysis revealed global microbial signatures that can make consistent prediction of colon cancer regardless of the cohorts, as well as cohort-specific microbial signatures that can shed light on confounders in colon cancer prediction. In this study, we also benchmarked the performance of different existing methods in making cross-cohort predictions, showing that StableMate is highly competitive. We noted that StableMate did not significantly outperform random forest, probably due to either a small number of cohorts affecting statistical power, or because of the difference between a linear logistic regression (StableMate) and a non-linear classification method (random forest). This, therefore, also motivated our simulation study where we considered a continuous response and generated enough repetition of experiments (Supplementary Figure S1, S2).

In the third case study we analysed scRNA-seq data of myeloid cells residing in the core and the surrounding periphery tissues of glioblastoma. We first identified a trajectory of continuous cell state transition between the cells at the two locations, then applied StableMate to identify stable and location-specifc genes associated with this cell state transition. By analysing periphery- and core-specific genes, we hypothesised that microglial polarisation seem to occur primarily in the tumour periphery, and the reprogramming of microglia into pro-tumour TAM happens after microglia infiltrate the tumour core. The stable genes exhibited consistent expression patterns in both the locations, hence ubiquitously involved in the development of both the pro- and the anti-inflammatory microglia.

In these case studies, the biological interpretation of the variable selections mainly focused on significant genes or microbial signatures. However, further experimental validations could hypothesise on the causal implication of stable predictors to the response.

StableMate is based on stabilised regression but implements a different algorithm for stochastic stepwise variable selection to select stable and environment-specific predictors with higher computational efficiency and accuracy. The stepwise framework of StableMate can be implemented with different base regressors to address different regression problems, such as ordinary least square regression and logistic regression, as we illustrated in our case studies. StableMate is available in R and can flexibly implement user-defined regression methods. One such extension could for example include non-linear regression methods, as well as penalised regression to avoid the pre-screening step currently proposed in StableMate.

## 4 Methods

### 4.1 Data and preprocessing

A summary of the data and StableMate analysis from the case studies is presented in Table 1.

#### 4.1.1 Breast cancer gene expression data

We analysed the RNA-seq dataset from The Cancer Genome Atlas Program (TCGA-BRCA) to study the transcriptional regulation of ESR1 in ER+ breast cancer (BC), available from the R package `TCGAbiolinks` (Colaprico et al., 2016). The dataset includes the expression quantification of 60,660 genes on 113 normal samples and 1,094 BC samples. We focused on the log-Transcript-Per-Million (logTPM) of 19,937 protein coding genes for analysis.

We used two other gene expression studies as validation: the microarray data of 2,509 BC samples from the METABRIC cohort (Curtis et al., 2012; Pereira et al., 2016), available from cBioProtal (Cerami et al., 2012) in the form of z-score relative to all samples (log), and RNA-seq data of 980 normal breast samples from GTEx (Lonsdale et al., 2013) in the form of logTPM.

#### 4.1.2 Colon cancer metagenomics data

We obtained nine CRC case-control studies of fecal metagenome from the R package `curatedMetagenomicData` (Pasolli et al., 2017). We excluded two studies with a sequencing depth lower than the average ten million reads per sample. The remaining studies included curated microbial species abundance and pathway abundance data from seven different countries and eight different cohorts: including 107 samples from Austria (Feng et al., 2015), 104 samples from the United States (Vogtmann et al., 2016), 125 samples from Germany (Wirbel et al., 2019), 509 samples from Japan (Yachida et al., 2019), 128 samples from China (Yu et al., 2017), and 114 samples from France (Zeller et al., 2014), as well as two cohorts containing 53 and 60 samples from Italy (Thomas et al., 2019). In total, all cohorts included 1,429 samples. We filtered the species and pathway abundance data from each cohort down to 313 species and 431 pathways that were detected across all cohorts.

To normalise the abundance data, we applied rank transformation by calculating the within-sample ranking quantile of the abundance of each species (or pathway). A species ranked the *q*^*th*^ most abundant in a sample is assigned the value (1 − *q*)/(*p* − 1), where *p* is the total number of species analysed. Therefore, the most abundant species of a sample has a rank transformed value of 1 and the least abundant species a value of 0.

#### 4.1.3 Glioblastoma single cell RNA-seq data

We analysed the glioblastoma (GBM) single cell RNA-seq (scRNA-seq) data from Darmanis et al. (2017) who sequenced single cells sampled from four GBM patients at their tumour cores and surrounding peripheral tissues. The raw and curated read count data included 3,589 cells measured on 23,368 genes available from http://gbmseq.org/. We retained 1,874 cells of myeloid cell types, including 1,329 cells sequenced from the core and 518 cells sequenced from the periphery for analysis. We used the R package `Seurat` to log normalise the data and identify the most variable 2,000 genes with the `FindVariableFeatures` function (Butler et al., 2018). We then imputed the log normalised data using Sincast imputation with default tuning (Deng et al., 2022) for StableMate variable selection and single cell projection. Diffusion map and diffusion pseudotime learning was performed on the original log-data (without imputation).

### 4.2 StableMate to identify stable and environment-specific statistical associations

We developed a variable selection method based on the stabilised regression framework proposed by Pfister et al. (2021), where the predictors and response are measured in different biological environments. The goal is to select *stable* and *environment-specific* (unstable) predictors that respectively make consistent and inconsistent predictions of the response across environments. A final model is built on the stable predictors, and is generalisable to unseen environments.

#### 4.2.1 The original stabilised regression

Briefly, stabilised regression (SR) examines all possible subsets of predictors in a brute force search, fits a regression function on each subset and evaluate the subset’s stability across environments and its prediction ability. First, the stability of predictor subsets are constructed based on either a Chow test (testing for equal regression coefficients of the predictors between regression functions fitted in a specific environment), or a resampling approach. Subsequently, prediction ability of stable subsets are evaluated based on negative mean squared prediction error combined with bootstrapping to define a cut-off for selecting the most predictive sets. The importance of each variable with respect to their stability, unstability, and prediction ability is then assessed via frequency of selection. The final SR model is obtained as a weighted average of the regression functions fitted on the stable and predictive subsets (refer to Pfister et al. 2021 for more details).

We identified several limitations of SR in its current form.

- It is computationally infeasible to enumerate every possible subset of predictors in ‘omics data where the number of predictors *P* is very large (i.e, > 30). Pfister et al. (2021) proposed the following solution: 1) pre-filter data to tens of predictors. Then from the pre-filtered predictor sets, 2) randomly sample thousands of subsets to test for stability and subsequently prediction ability. However, we argue that this solution is inefficient, as thousands of subsets is not sufficient to represent the subset space of many predictors. A drastic pre-filtering is therefore required but can result in filtering out important predictors.
- Identifying first the stable predictor sets, then assess their prediction ability is not efficient. This not only because the stable and predictive sets are included in the predictive sets, as we describe in Supplemental Methods 7.1.1, but also because stability is more difficult to compute compared to prediction ability.

Because of these limitations, SR results lack both variable selection and prediction accuracy for large datasets, as we highlight in our simulation (Supplemental Figure S1).

##### The StableMate approach

StableMate addresses these issues by 1) implementing a greedy rather than a brute force approach to select predictor sets based on an improved version of stochastic stepwise regression (ST2*), which is a stochastic selector, 2) building a variable selection ensemble using repeated ST2*, 3) pre-screening predictors before each ST2* using random Lasso to enable a much larger starting set of predictors than SR, 4) identifying first the predictive variables, then narrowing down to the stable predictors to be more efficient in the search, 4) developing the concept of pseudo-predictor to benchmark ST2* selections. The full methodological details are available in Supplementary Methods 7.1.3.

#### 4.2.2 Main steps of StableMate

We summarise the main steps of StableMate, a more detailed algorithm is presented in Algorithm 1 in Supplemental Methods 7.1.3.

1. Depending on the type of analysis, a base regressor for ST2* is first specified, for example, we used Ordinary Least Square regression for case studies in Sections 2.2 and 2.4 and Generalised Linear Models in Section 2.3.
2. For each iteration *k, k* = 1, …, *K*
  a. Apply random Lasso pre-screening, then add pseudo-predictor
  b. (Run ST2* to select the most predictive predictor set denoted 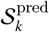.
  c. (Run ST2*to select the stable predictors within 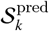 such that 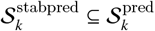 .
3. Define importance score for prediction and stability.
4. Calculate significance cut-off scores to define stable, unstable and non-significant variables.
5. Fit the final ensemble regression model as weighted average of the fitted regressions on 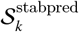

#### 4.2.3 Pre-screening predictors based on random Lasso

We first pre-filter predictors based on a random Lasso procedure. For each ST2* run (described below), we randomly sample one half of samples to select the top *p* predictors with Lasso (Tibshirani, 1996). As an example, we chose *p* = 100 for Sections 2.2 and 2.4. The advantages are of two folds. First, across the different resampling runs, the top *p* predictors are expected to differ thus enabling to cover a large and diverse range of predictors in our overall search. Second, we improve the stability of Lasso pre-screening when subjected to sample perturbation (Meinshausen and Bühlmann, 2010) (see more details in Supplementary Methods 7.1.6).

#### 4.2.4 ST2*: a new stochastic stepwise variable selection procedure

We improved the ST2 algorithm proposed by Xin and Zhu (2012). ST2 is a stochastic version of the classic stepwise variable selection. ST2 searches for a set of predictor maximising a particular objective function to quantify the predictive ability or stability of predictor sets. It uses a greedy approach with iterative forward and backward searching steps. ST2 starts with an initial predictor set (which can be empty). In the forward step, a collection of predictor sets are randomly sampled from predictors that are not included in the current model. The predictor set which yields the largest increase of the objective function is then added to the current model. In the backward step, a collection of predictor sets is randomly sampled from predictors that are included the current model. The set which yields the largest increase of the objective function is then removed from the current model. The forward and backward steps alternate until the objective function does not improve further. However, the major drawback of ST2 is that it samples subsets of a fixed size that is randomised at each step. If a wrong size is sampled, ST2 may stop prematurely, leading to inaccurate variable selection.

In ST2*, we follow the ST2 algorithmic framework but we improved the procedure to sample different predictor subset sizes at each step (refer to the Supplementary Methods 7.1.4 for a detailed description of the ST2* algorithm.). We also added objective functions that are well suited to assess prediction, and stability, namely, we used the Negative Bayesian Information Criterion (NBIC) and Negative Prediction Sum of Squares (NPSS) (Supplementary Methods 7.1.5).

Finally, we run ST2* on *K* iterations (for example *K* = 2000 in our case studies) first to identify the predictive subsets of predictors, second to identify the stable subsets within each predictive subset. As a result, we select an ensemble of stable and predictive predictor sets. These iterations address the stochastic nature of ST2* which can yield to potentially different predictor sets for each iteration. The sets of stable predictors are then used to build the final regression model (described below).

#### 4.2.5 Cut-off prediction and stability scores

We calculate a prediction score of each predictor based on how often the predictor is selected as predictive across the ensembles. We do similarly for the stability score. The output can be represented in a variable selection plot such as Figure 2A, where the scores are represented on the *x*-axis (prediction) and *y*-axis (stability).

To define a significance cut-off of these scores, we create a pseudo-predictor as a negative control. A pseudo-predictor is represented as an artificial index *P* + 1 so that its inclusion in the regression model does not affect the model fitting nor the value of the objective function in ST2*, but it is still taken into account when calculating the scores of all predictor sets.

We applied a bootstrap procedure on the variable selections to compare the distributions of the prediction and stability scores of the predictors to that of the pseudo-predictor to assess their significance. A predictor with a prediction score larger than the pseudo-predictor’s in more than 97.5% times of the bootstrap iterations is considered as significantly predictive. We do similarly for the cut-off stability score. A predictor is considered environment-specific (unstable) it its stability score is lower than that of the pseudo-predictor. Finally, the rest of the predictors is assigned as ‘Non-significant’, as shown in plot Figure 2A. Note that since these cut-off scores are based on bootstrap of variable selections the significance indicates the variability in the ST2* selections, and hence provide a reference on whether more ST2* runs need to be performed.

#### 4.2.6 Final ensemble regression model generalisable to unseen environments

The final regression model is then built on the different sets of stable and predictive predictors. Each regression model is fitted by regressing the response variable on each stable and predictive subset in the ensemble. We then aggregate these models as the average of the fitted regression weighted by the ranking of objective functions NBIC and NPSS.

### 4.3 Principal component analysis

We used the *prcomp* function from the R package `stats` (R Core Team, 2013) to perform Principal Component Analysis (PCA).

#### Gene modules

PCA (centered but not scaled) was used to identify meta-genes in the form of principal components that represent gene modules from the TCGA breast cancer RNA-seq data. The meta-genes were then used as the predictors of ESR1 expression for the subsequent StableMate analysis. To avoid overfitting, ESR1 was removed from the data.

#### Aggregation of gene expression with similar expression patterns

We applied PCA (not centered nor scaled) on the set of genes, and extracted the loading coefficients of each gene on the first principal component using a soft-thresholding approach to identify the top contributing genes with loading coefficients of the same sign. We then considered the absolute value of the loading coefficients of these top genes to obtain positive weights, which we then used for aggregating their expression by a linear combination.

### 4.4 Principal coordinates analysis

We used the *cmdscale* function from the R package `stats` (R Core Team, 2013) to perform Principal Coordinates Analysis (PCoA). PCoA was performed on the combined colon cancer case-control studies with classical multidimensional scaling on Euclidean distances between samples. We calculated two distances matrices on either the 313 species of the full data and on the 23 species selected by StableMate. Permutational Multivariate Analysis of Variance (PERMANOVA) was then used to test the separation of the sample groups based on disease status, or cohorts. We used the *adonis* function from the R package `vegan` (Oksanen et al., 2022).

### 4.5 Methods benchmark

We benchmarked the prediction performance of StableMate against other commonly used methods, including Ordinary Least Square regression (OLS), Generalized Linear Model (GLM or logistic regression), Lasso regression (Lasso, Tibshirani (1996)) and Random Forest (RF, Breiman (2001)).

The regression models were trained for the different benchmarking tasks described below on the pooled data of training environments. For predicting continuous responses in the simulation study, we used GLM Lasso with a Gaussian family. For binary classification of colon cancer in the second case study, we used GLM Lasso with a binomial family. StableMate requires to specify the different sample environments, while in RF samples were weighted according to the inverse of the size of the environment each sample belongs to. Lasso penalties were tuned using cross-validation, where each environment is used as a fold to minimize the averaged mean squared error. We used the functions *lm* for OLS, *glm* for GLM, *cv*.*glmnet* from the package `caret` for Lasso (Kuhn, 2022) and the R package `randomForest` for RF (Liaw and Wiener, 2002).

#### Simulation study

The benchmark results are shown in Supplementary Figures S2 and S1.

We simulated systems of variables observed from four environments. A system is a model that describes the causal relationships between variables, and an environment is the probability distribution of variables that generates data. Therefore, a system of variables in different environments are generated by different probability distributions but with the same causal relationships. The simulations are described in Supplementary Methods 7.2.1. For each simulation run, a variable in the system was randomly sampled as the response while the remaining variables were set as predictors. The regression models were trained on data generated in the first three environments to predict the response. The data of the fourth environment was used for testing.

#### Metagenomics case study (Section 2.3)

We trained each regression model to predict colon cancer disease status. We performed Leave One Dataset Out (LODO) cross-validation. We considered either all 313 species, or 431 pathway abundances (no pre-filtering). In addition, we also performed LODO validation to evaluate the performance of the RF models trained using the different sets of predictors selected by either StableMate, Lasso and RF.

### 4.6 Diffusion map and diffusion pseudotime

#### Diffusion map

In case study 3 (Section 2.4) we visualised the scRNA-seq data using Diffusion Map (DM), which is a non-linear dimension reduction method highly suitable for single cell data with potential cell state transitions. DM learns transition probabilities between cells and projects cells into a lower dimensional Euclidean space that approximate the ‘diffusion distances’ between cells accordingly. DM was run on the most 2,000 most variable genes of the log normalised data using the R function *DiffusionMap* with default parameters from the package `destiny` (Haghverdi et al., 2015).

#### Diffusion pseudotime

Diffusion pseudotime (DPT) inference was then applied following DM learning (Haghverdi et al., 2016). The cell that has the largest DC1 score in the tumour periphery was chosen as the root of the cell trajectory. (Figure S8B). The distance between the cumulative transition probabilities of any cell with the root cell is defined as its pseudotime.

### 4.7 Sincast projection of scRNA-seq onto a reference atlas of myeloid cells

We used Sincast (Deng et al., 2022) available at https://github.com/meiosis97/Sincast to impute scRNA-seq data and to query GBM cell types and cell states. We queried the identity of a specific subset of GBM cells, namely myeloid cells classified by Darmanis et al. (2017). The reference myeloid atlas was from Rajab et al. (2021) who compiled bulk RNA-seq and microarray data of myeloid cells from 44 independent studies. Sincast projects the query scRNA-seq cells onto the atlas by calculating predicted principal components of the cells, that are then represented on the PCA of the reference atlas. The result is a 3D PCA plot (i.e., Figure 4A) where we can infer the identity of the query cells according to the biology of their surrounding atlas samples.

#### Sincast imputation

The log normalised scRNA-seq data were imputed using the *sincastImp* function with its default parameters, where the imputation of any cell is cased on its nearest neighbouring cells.

#### Sincast projection

Only the most 2,000 most variable genes of the query scRNA-seq data were considered. The query data were projected onto the reference PCA atlas after rank transformation. The projection is then reorganised and visualised via diffusion map.

#### Quantification of query cell identity

To quantify the identity of each query cell based on the reference cell types, we used Sincast modified version of Capybara cell scores of Kong et al. (2022). The approach is based on Weighted Restricted Least Square regression (Deng et al., 2022).

## Declarations

### Code availability

The StableMate R code and data analysis are available at https://github.com/meiosis97/StableMate.

## Acknowledgements

We thank Miguel Ángel Berrocal Rubio and Patrick Lichtner for their helpful insights regarding the variable selection results from the GBM case study.

## 5 Supplementary Results

### 5.1 CRC metagenomics meta-analysis: low fiber and high protein diet can confound colon cancer prediction in the Austrian cohort

In traditional meta-analysis, we can identify cohort-specific species predictive on CRC in one cohort but not in the others. However, this type of analysis limits comparisons of the selected species across cohorts. In contrast, StableMate defines cohort-specificity in a more comprehensive way by incorporating stability as a measure that quantifies the consistency of CRC-species association pattern across cohorts.

As an example of Austrian-specific species with StableMate analysis, we identified *Prevotella copri* and *Bacteroides xylanisolvens*, that are known to be associated with high-fiber non-Westernized diet and function in fiber digestion (Figure 3C) (Despres et al., 2016; Yeoh et al., 2022). Notably, *Prevotella copri* was the most predictive species but with a stability score close to zero. In the Austria cohort, these two species were merely detected in normal samples but were dominant in the CRC samples (Supplementary Figure S4). Interestingly, this pattern was not observed in the other cohorts, especially In the Asian cohorts (Chinese and Japanese cohorts), these two species were abundant in normal samples. Our observations were consistent with the common dietary habits of the countries of the cohorts.

To further gain functional evidence on dietary effects on CRC prediction, we queried Austrian-specific pathways using StableMate and identified the L-arginine biosynthesis III pathway that is enriched in the Austrian CRC patients but not in the other cohorts (Supplementary Figure S6A,D). The abundance of this pathway has been found to be potentially related to the intake of protein-rich diet as an important source of L-arginine in human body other than self-production. The Austrian diet is protein-rich, which could explain the low L-arginine synthesis activity observed in the healthy individuals of the Austrian cohort (Supplementary Figure S6C).

Therefore, StableMate enables us to formulate a plausible explanation of the specificity in the Austrian cohort, which may be attributed, in part, to the population’s relatively low daily consumption of dietary fiber, as well as an increased fiber intake subsequent to a diagnosis of CRC.

## 6 Supplementary Figures

**Figure S1.**
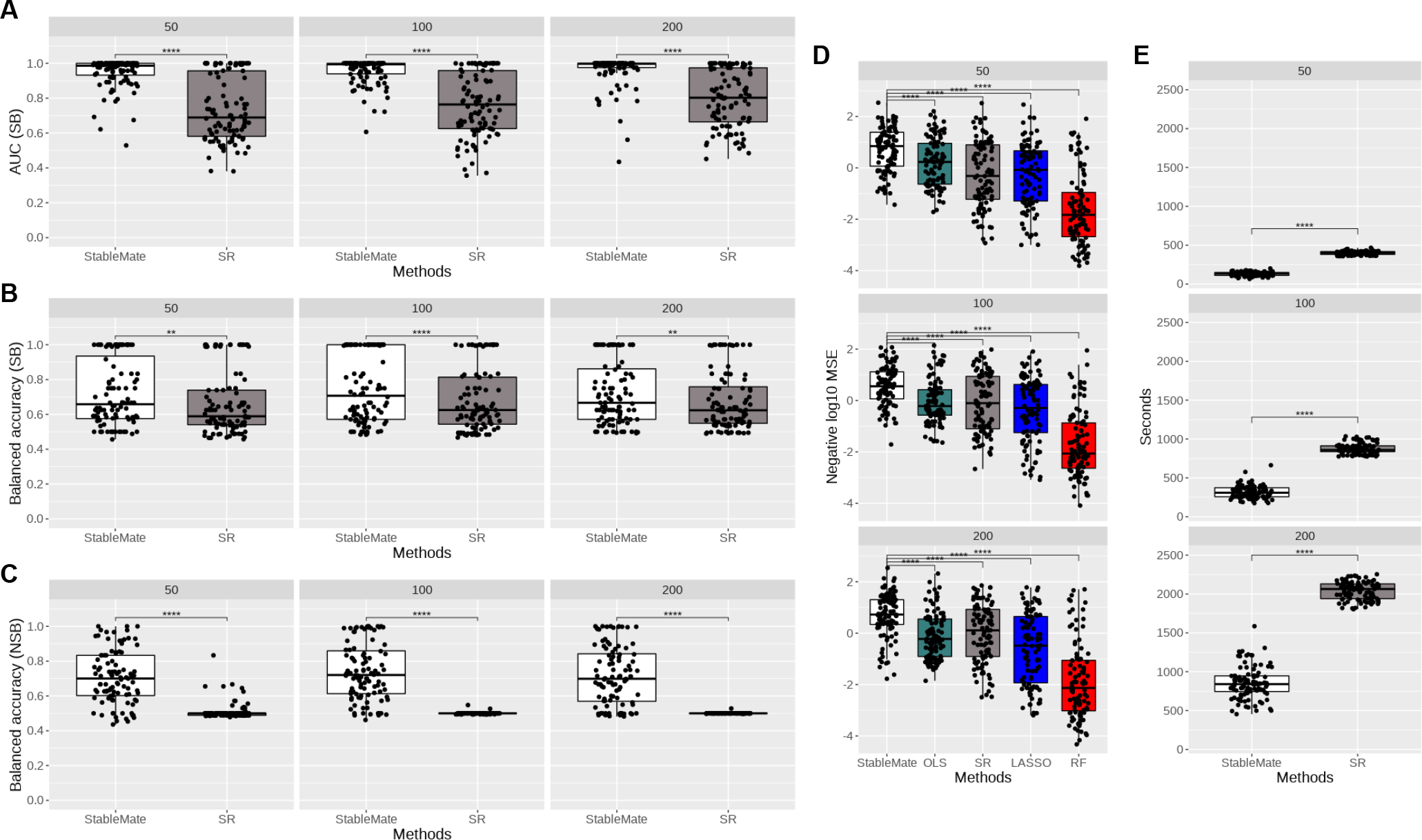
Benchmark of StableMate in selecting truly stable and unstable predictors and in building generalisable prediction models. To benchmark StableMate against stability regression (SR) in selecting truly stable and unstable predictors, we simulated datasets with 50, 100 and 200 predictors, where the subsets of truly stable and unstable predictors were known. For each choice of the number of predictors, 100 datasets were simulated from a structural causal model, each comprising equal number of samples from four environments (see Supplementary Methods 7.2.3). In all sub-plots, p-values of paired t-tests between StableMate and one of the competing methods are represented by the number of asterisks ∗ (∗ : *p* < 0.05, ∗∗ : *p* < 0.005 etc.). **(A)** Area under the curve (AUC) values for correctly selecting the stable predictors. According to the paired t-test results, StableMate had significantly higher AUC values in all cases. **(B)** and **(C)**: balanced accuracy, i.e. average of sensitivity and specificity, in selecting stable and unstable predictors. According to the paired t-test results, StableMate had significantly higher balanced accuracy in all cases. To benchmark StableMate against the ordinary least squares (OLS) regression, SR, the Lasso regression and random forest (RF) in building generalisable prediction models, we used the same simulated data. We trained the regression model on three out of the four environments and tested on the fourth environment. **(D)** Negative mean squared error for predicting in testing environments. According to the paired t-test results, StableMate outperformed all competing methods with a significantly higher accuracy. **(E)** Computation times. StableMate was at least two times faster to compute compared to SR.

**Figure S2.**
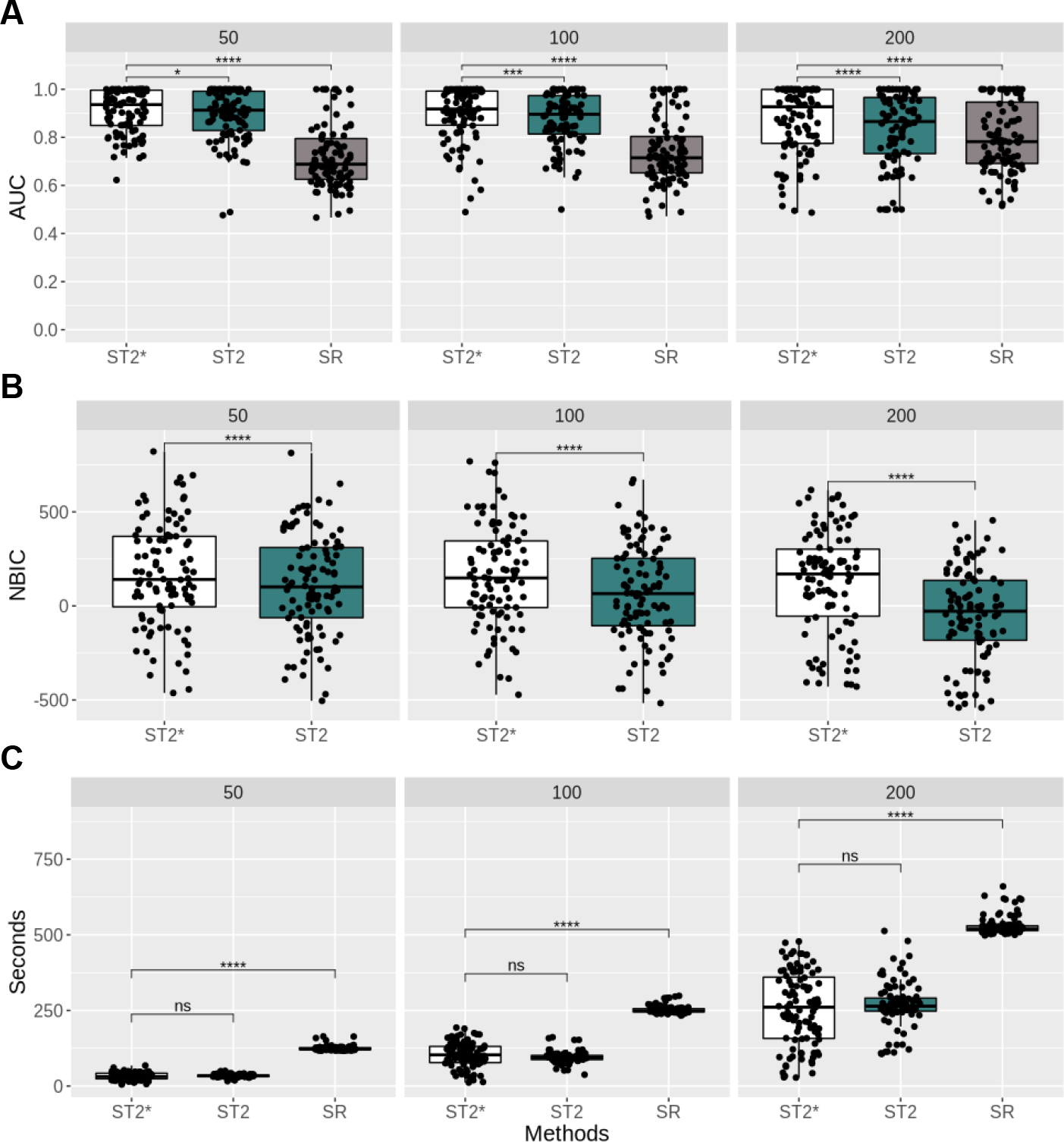
Benchmark of our proposed ST2* in selecting truly predictive variables, against stabilised regression (SR) and stochastic stepwise selection (ST2). We simulated datasets with 50, 100 and 200 predictors, corresponding to the three columns above, where the subsets of truly predictive variables were known. For each choice of the number of predictors, 100 datasets were simulated according to a structural causal model (see Supplementary Methods 7.2.2 for details). For each dataset, ST2 and ST2* were run 100 times to build variable selection ensembles. In all subplots, p-values of paired t-tests between ST2* and one of the competing methods are represented by the number of asterisks ∗ (∗ : *p* < 0.05, ∗∗ : *p* < 0.005 etc.). **(A)** Area under the curve (AUC) values for correctly identifying the predictive variables. Although the difference between ST2E* and ST2E was not large in terms of the absolute values of AUC, the paired t-test results suggested that the AUC values of our ST2E* were significantly higher the other two methods in all three cases. In particular, the superiority of ST2* over ST2 became increasingly significant when the total number of predictors increased. **(B)** Mean negative Bayesian information criterion (NBIC) across the ensemble of ST2* and ST2 for the further comparison of ST2* and ST2. Again, ST2* outperformed ST2 in all cases by having significantly higher NBIC values. **(C)** Computation time in seconds. ST2* and ST2 had comparable computation times. Both methods were significantly faster than SR.

**Figure S3.**
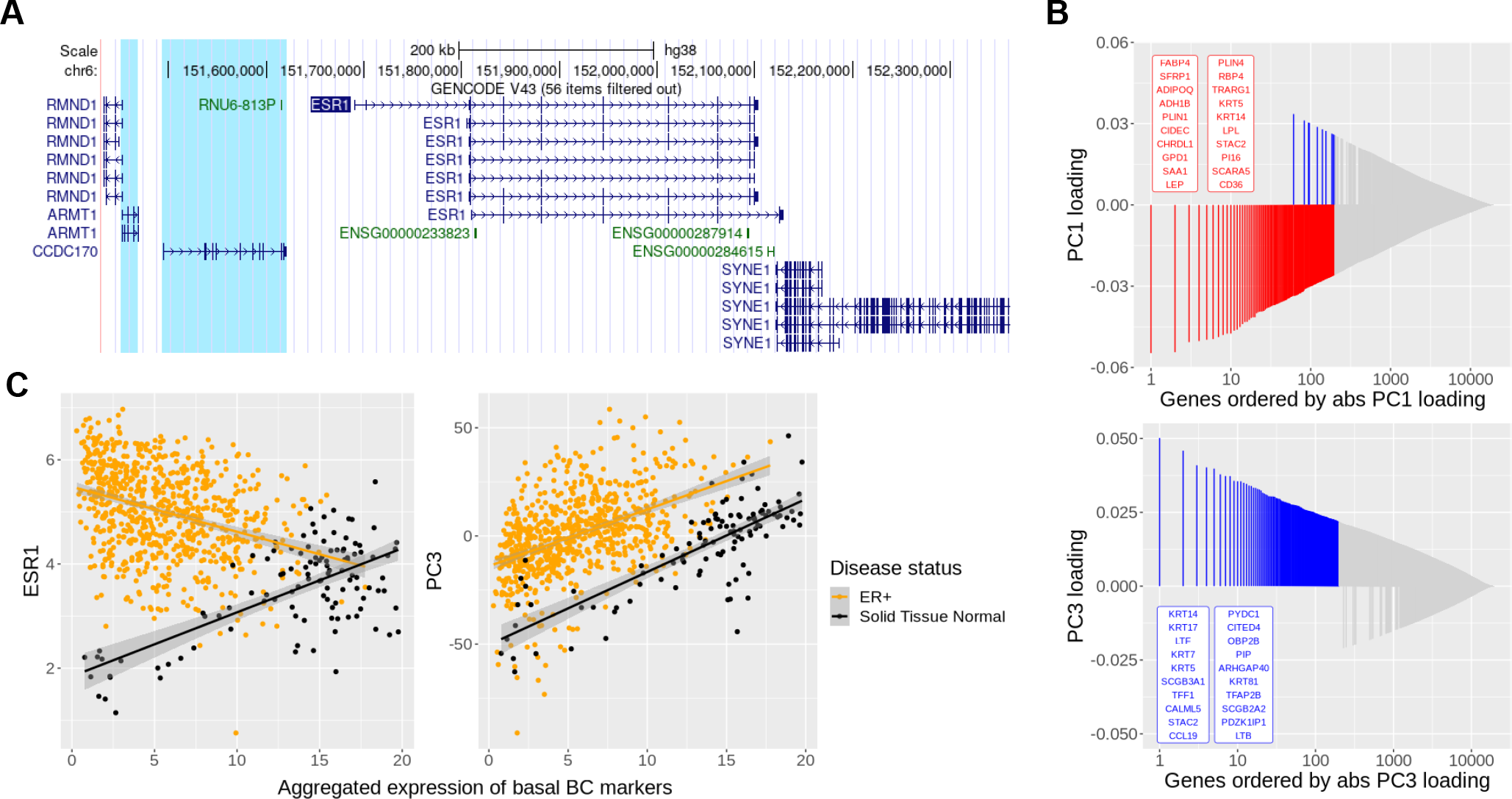
Breast cancer case study (Section 2.2). **(A)** UCSC Genome Browser view of ESR1 annotated by GENCODE v43. StableMate identified ARMT1 and CCDC170 as stable. These two genes are the closest to ESR1 in the upstream genomic region of ESR1 and are subject to similar transcriptional regulation as ESR1. This may explain why there were identified as stable predictors by StableMate. **(B)** Loading plots of the first and the third principal components (PC1 and PC3) shown in Figure 2C. Genes were ordered, in descending order, by the absolute value of their loading coefficients in each component (*x*-axis), and the top 200 genes are coloured according to their sign (positive in blue and negative in red). The names of the top 20 genes are shown. Genes dominating PC3 included multiple basal cytokeratins with positive loading coefficients, suggesting that PC3 is positively correlated with basal cytokeratin expression. **(C)** ESR1 expression (left panel, similar to Figure 2C) and PC3 score (right panel) against aggregated expression levels of basal BC signature genes, which were retrieved from Li et al. (2022). The expression level of basal genes were positively correlated with ESR1 expression in normal samples and negatively in ER+ breast cancer. Recall that we observed similar patterns between PC3 score and ESR1 expression in Figure 2C, suggesting that PC3 was associated with ‘basalness’ characteristics of the samples. The positive correlation between PC3 and basal genes in the right panel further validated our hypothesis.

**Figure S4.**
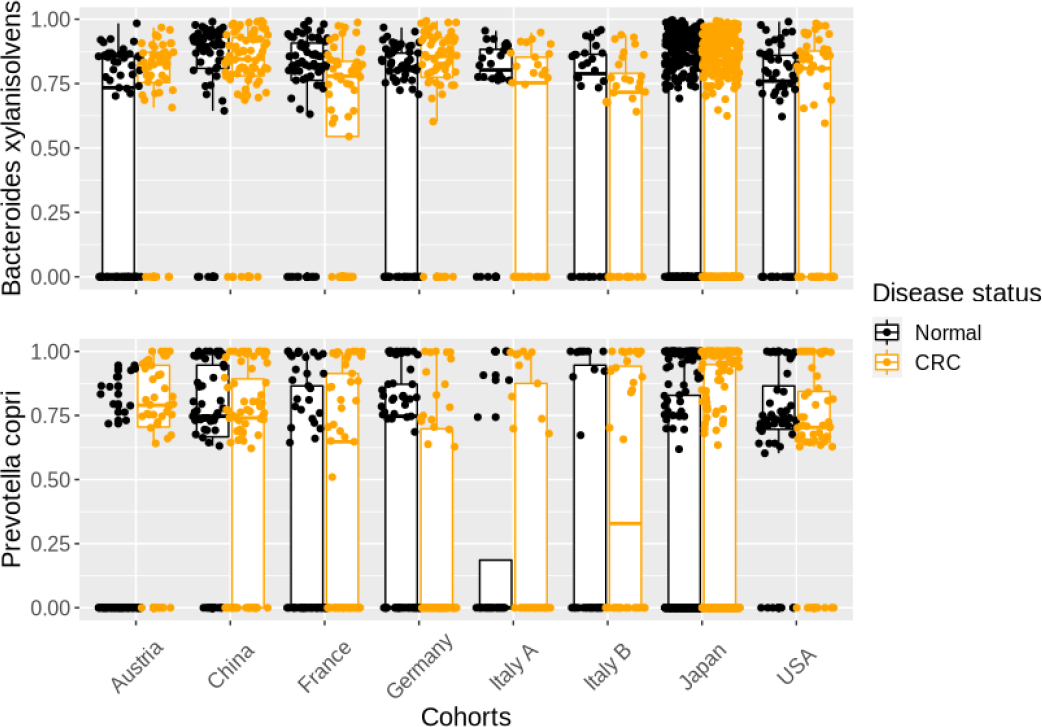
Colon cancer metagenomics case study (Section 2.3). Boxplot showing the abundance of *Prevotella copri* and *Bacteroides xylanisolvens* that were selected as Austrian-specific. These two species are known to relate to fiber digestion, showing enrichment in the CRC samples of the Austrian cohort, but not in the Asian cohorts.

**Figure S5.**
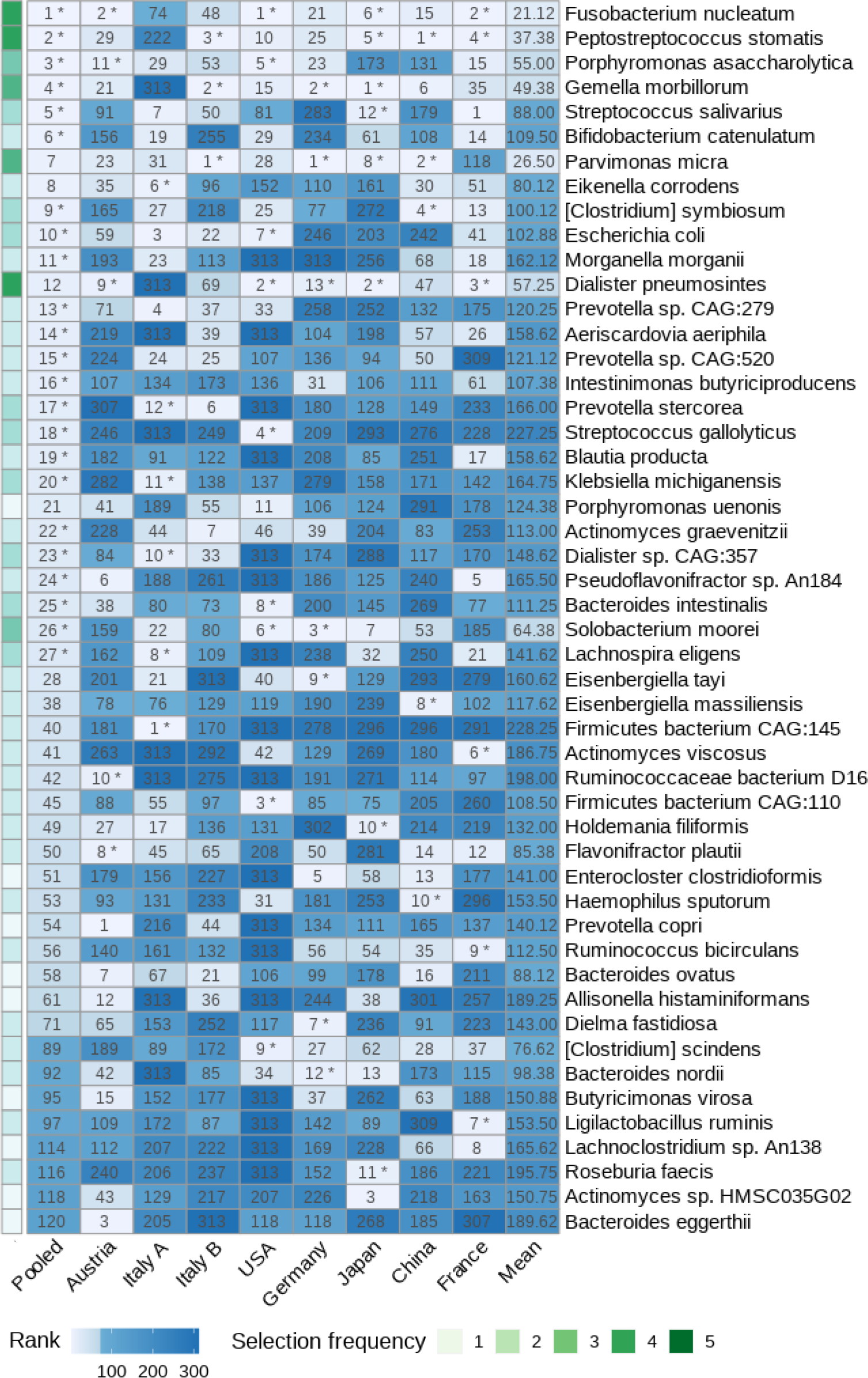
Colon cancer metagenomics case study (Section 2.3). Prediction scores of the selected species predictive of CRC (rows). StableMate was applied to select species predictive of CRC on all eight cohorts/countries combined, refered as ‘*Pooled*’ in the first column, or on each separate cohort in the other columns. The rank of the species selected as predictive is shown (low rank = high prediction). The last column (‘*Mean*’) averages the ranks across all cohorts. Rows are ordered according to the Pooled column ranks. The stability score was calculated by setting cohorts and the environment variable. Asterisks indicate stable species whose prediction on CRC is consistent regardless of the cohort. The frequency of stability selection across each of the eight cohorts is indicated, showing that species with high prediction scores in the pooled analysis were mostly stable and more frequently selected as stable across individual cohorts.

**Figure S6.**
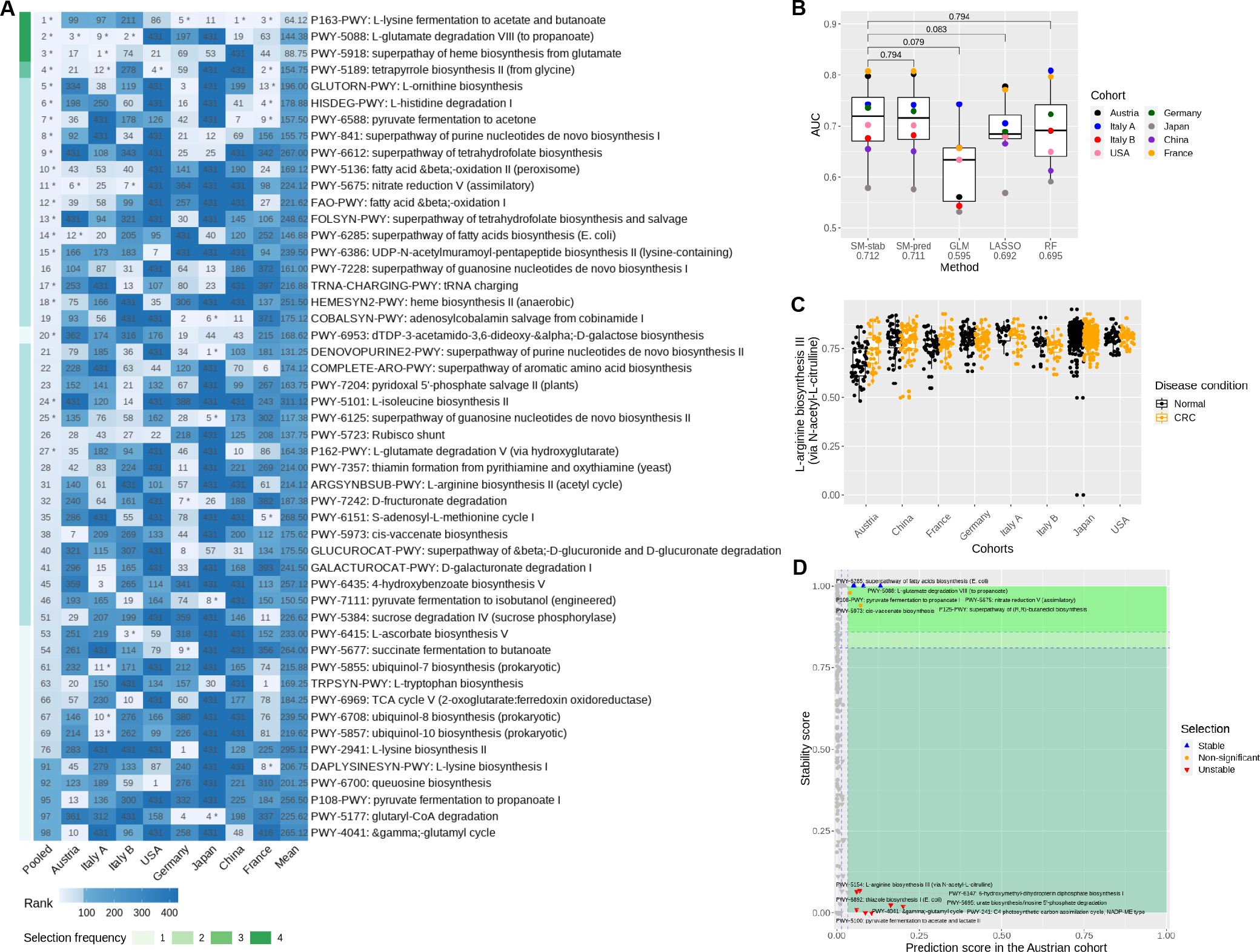
Colon cancer metagenomics case study (Section 2.3) on the pathway abundance data. **(A)** StableMate selects metabolic pathways predictive of CRC. Annotation and interpretation of this figure is similar to Figure 3B but indicate pathway names. Compared to Figure 3B, the prediction scores in the individual cohorts were less consistent. **(B)** Benchmark of prediction performance with Leave One Dataset Out (LODO) cross validation, similar to Figure 3C. StableMate outperformed the other methods with the highest mean AUC scores. **(C)** Boxplot showing the abundance of L-arginine biosynthesis III pathway, which was selected as predictive in the Austrian cohort, but was identified as unstable. L-arginine, which can either be synthesised by human body or be taken from high-protein diet, was enriched in the CRC samples of the Austrian cohort, but not in the other cohorts. **(D)** StableMate selection on CRC predictive pathway in the Austrian cohort only.

**Figure S7.**
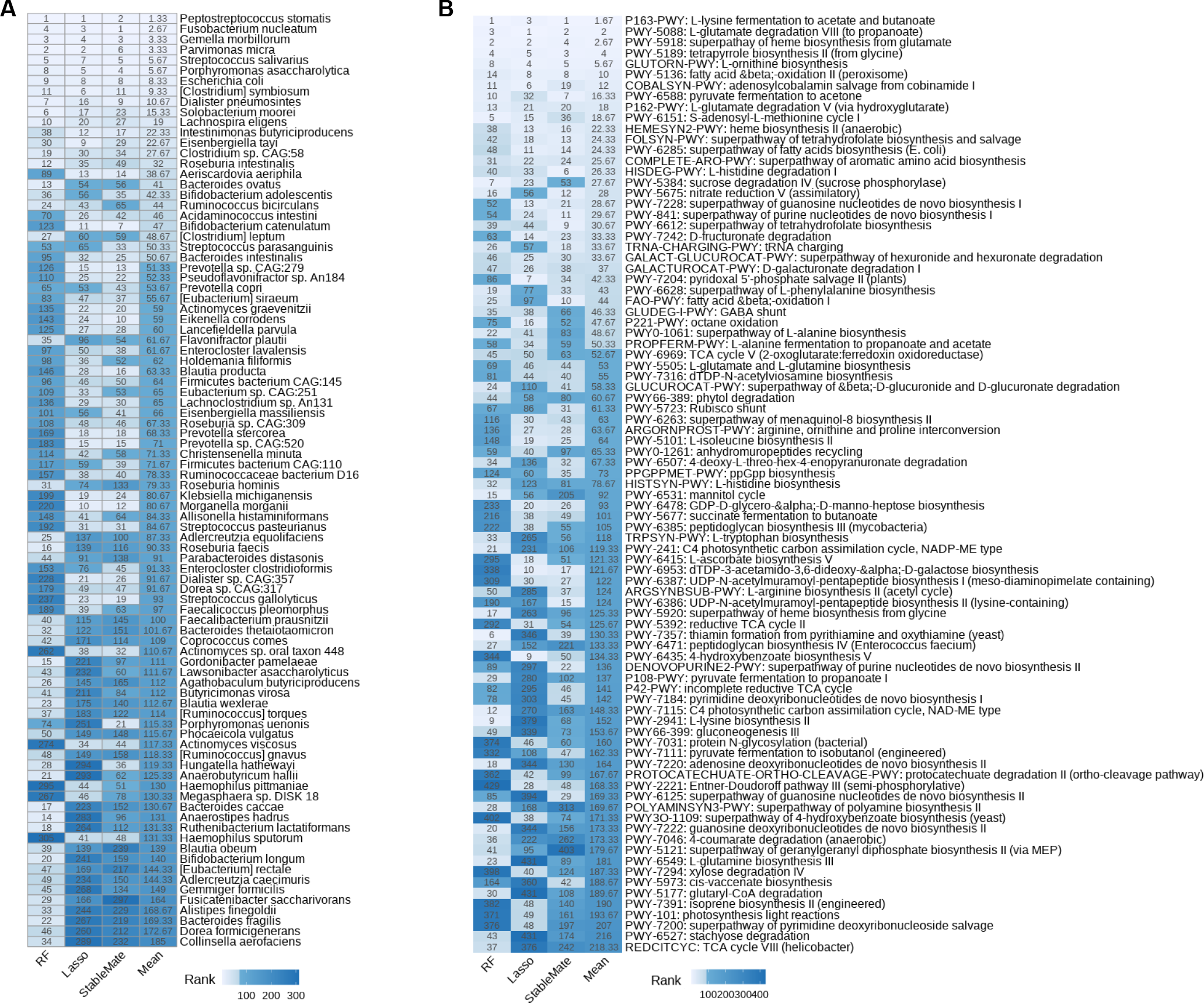
Colon cancer metagenomics case study (Section 2.3): comparisons between StableMate, Lasso and Random Forest (pooled data of all cohorts). **(A)** The top 50 predictive species were selected by either StableMate, Lasso and RF and visualised in a heatmap where cells are colored and labeled according to their selection ranks. Selected predictors were ranked according to stability and prediction scores for StableMate; their order in entering the Lasso selection path for Lasso; importance scores for RF. Rows represent species and columns represent methods, with the last column (labeled by ‘Mean’) depicts the average ranking across the three methods. Rank 1 indicates that a species is selected as top predictor. **(B)** Similar to **(A)** for the pathway abundance data, showing a lower consistency between methods compared to the species abundance data.

**Figure S8.**
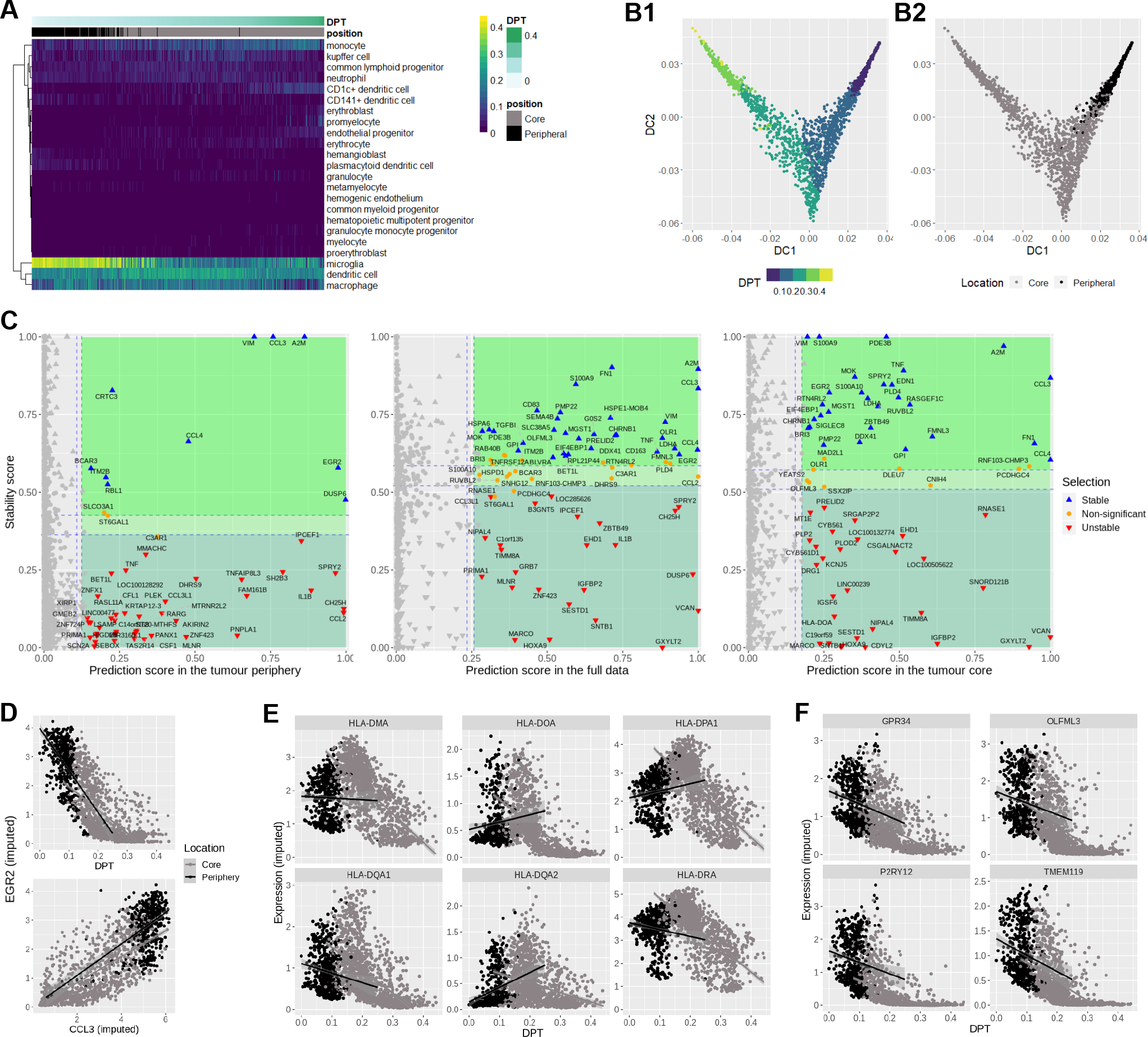
Glioblastoma scRNA-seq case study (section 2.4). **(A)** Heatmap illustrating Capybara quantification of cell identity based on a reference myeloid cell atlas (Rajab et al., 2021). Columns represent query cells and were ordered from left to right by increasing DPT. Rows represent reference myeloid cell types. Peripheral cells, which were located mainly at the start of the inferred trajectory, obtained high scores on microglia. The identity of core cells were heterogeneously shared among monocytes, dendritic cells, macrophages and microglia. **(B)** Data visualization using diffusion map, showing a continuous cell state transition from the cells at the tumour pheriphery. **(B2)** correspond to (B1) and (A2) respectively, but cells are colored according to Diffusion Pseudotime (DPT), illustrating a cell state transition. **(C)** Genes selected as predictive of pseudo-time (DPT) by StableMate – illustrated in Figure 4A, where cells were pooled from both location (left panel), or were only from the tumour periphery (middle panel) or the tumour core (right panel). Genes that were stable in the full data but unstable in either location, such as TNF, were considered false discovery. **(D)** DPT (top) and the expression of stable cytokines CCL3 (bottom) versus the expression of EGR2 imputed with Sincast. EGR2 is selected as a stable predictor for both DPT and CCL3 by StableMate. **(E)** Expression levels of the genes that encode alpha-chains of MHC-II molecules, including HLA-DOA, which was selected as one of the core-specific genes (imputed by Sincast). The MHC-II genes displayed no correlation with DPT in the periphery and strong negative correlation with DPT in the core; **(F)** Expression levels of microglia markers identified by Darmanis et al. (2017) imputed by Sincast. The expression pattern of these genes was similar to that of HLA-DOA but with a large decrease in expression levels in the core.

**Figure S9.**
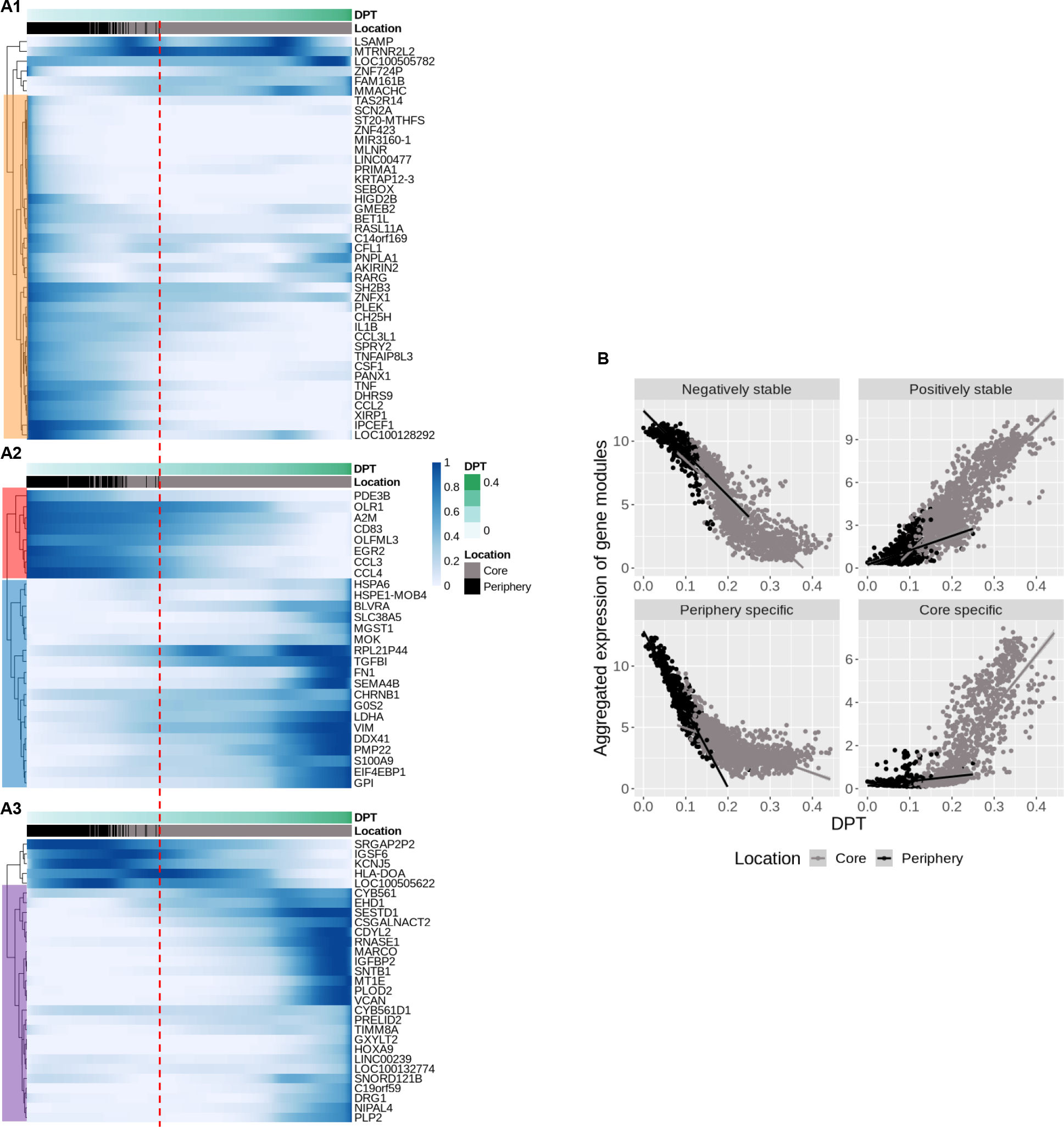
Glioblastoma scRNA-seq case study (section 2.4). **(A)** StableMate was applied to select genes predictive of DPT, where cell location (core and periphery) was set as the environmental variable. Four gene modules were identified via hierarchical clustering. In the heatmap of the min-max transformed gene expression levels, cells in columns are ordered according to pseudo-time, and genes in rows are only labelled if mentioned in the text. Left margin highlights the major gene modules of the periphery-specific genes (tumor periphery predictive genes, **(A1)**), the stable genes (consistently associated with DPT, **(A2)**) and the core-specific genes (tumor core predictive genes, **(A3)**). The red dashed line indicates the DPT of the last peripheral cell. The stable genes in A2 can be clustered into two major groups based on their correlation sign between their expression levels and DPT. The expression levels of the periphery-specific genes were mostly negatively correlated with DPT in A1, whereas the expression levels of the core-specific genes were mostly positively correlated with DPT in A3. **(B)** Aggregated expression levels of the four predictive gene modules identified in (A). Genes were either ‘negatively stable’ (stable and negatively correlated with DPT), ‘positively stable’ (stable and positively correlated with DPT), periphery-specific or core-specific.

## 7 Supplementary Methods

### 7.1 StableMate: stabilised regression to identify stable and environment-specific predictors

Stabilised Regression (SR) is a variable selection framework aiming at finding predictors that are predictive of the response variables and stable across multiple environments, such that the resulted regression model is generalisable to unseen environments (Pfister et al., 2021). In system biology, an environment can refer to either a technical or biological condition, such as batch, disease state or patient, under which the data are generated. It is also of interest to identify unstable predictors that are predictive of the response, but their relationship with the response is not stable across environments. We introduce StableMate, an SR framework which (1) selects and distinguishes predictors with high predictive ability that are either stable or unstable; and (2) builds a regression model based on the stable predictors that is generalisable to unseen environments.

#### 7.1.1 Mathematical setting

Consider a set of environments ℰ. In each environment *e* ∈ ℰ, observations are distributed as (*X* ^*e*^, *Y* ^*e*^), where 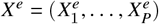 denotes the *P* predictors and *Y* ^*e*^ denotes the response.

We assume that, for each *e* ∈ ℰ, there exists a subset 𝒮 ⊆ {1, …, *P*} such that

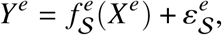

where 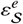 is a zero-mean noise term and where 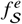 represents the true relationship between the response *Y* ^*e*^ and the predictors 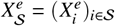. Note that 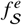 does not depend on predictors outside 𝒮 and hence 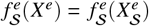.

The goal of SR is to find the stable and predictive predictors of *Y* . Such predictors are defined as members of the *Stable Blanket* (**SB**), denoted as 𝒮^sb^, defined as the smallest subset 𝒮 ⊆ {1, …, *P*} of predictors that satisfy the following conditions. They are both

1. Generalisable: for any environments (*e, e*^′^) ∈ ℰ, we have 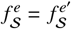 ; and
2. Regression optimal with respect to all generalisable sets: 𝒮 achieves the lowest pooled mean square error among all generalisable sets, such that, for all predictor set 𝒮^′^ ⊆ {1, …, *P*} satisfying the generalisable condition 1 above, we have

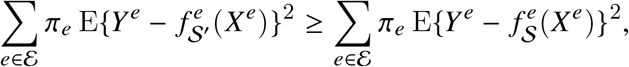

where *π*^*e*^ denotes the probability that an observed sample belongs to environment *e*.

The generalisable condition 1 requires that we use the same regression function 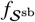, independent of *e*, to describe the relationship between *Y* ^*e*^ and *X* ^*e*^ across all environments (*e* ∈ ℰ). The regression optimal condition 2 requires that regressing *Y* ^*e*^ on predictor set 𝒮^sb^ yields the best prediction across environments. The predictor set satisfying both conditions may not be unique, hence we define 𝒮^sb^ as the smallest of such sets. The reader can refer to Pfister et al. (2021) for their discussion on identifiability of SB and their relation to causal inference.

We define the *Markov blanket* (**MB**), denoted as 𝒮^mb^ ⊆ {1, …, *P*}. MB is the smallest subset of predictors that is regression optimal with respect to *all predictor sets*, whereas SB is regression optimal with respect to *all generalisable sets*. Hence SB is always a subset of MB, and the difference of the sets 𝒮^mb^ − 𝒮^sb^ is referred to as the *non-stable blanket* (**NSB**). NSB predictors are predictive but not stable. They are useful to identify predictors whose relationships with the response change across environments. In the main text, for simplicity, we refer to predictors in SB as ‘stable predictors’ and those in NSB as ‘environment-specific predictors’.

#### 7.1.3 The original SR algorithm

The goal of SR is two-fold: to identify SB and NSB, and to build a regression model that is generalisable to unseen environments with SB. Motivated by the fact that SB is regression optimal among generalisable sets, SR selects SB by identifying all generalisable sets first and assemble the most predictive (regression optimal) ones to build a variable selection ensemble. An outline of the SR algorithm is given below:

1. Conduct a statistical test for the generalisable condition on each predictor subsets 𝒮 ⊆ {1, …, *P*}. Build an ensemble of generalisable subsets ℰ^gen^ that pass the test.
2. 2.Conduct a statistical test for the regression optimal condition on each generalisable subsets 𝒮 ∈ ℰ^gen^. Build an ensemble of generalisable and regression optimal subsets ℰ^genopt^ that pass the test.
3. 3.An importance score (e.g, selection frequency) is calculated based on ℰ^genopt^, for *p* = 1, …, *P*, to evaluate how stable and predictive each predictor *p* is in terms of how well it satisfies the generalisable and regression optimal conditions. A final estimate of SB is generated by selecting predictors with the highest importance score.
4. 4.A regression model is fitted using each 𝒮 ∈ ℰ^genopt^. All models are aggregated (e.g., by taking the average) into a final model for prediction.
5. 5.Repeat step 1 to 3 multiple times, each time on a re-sample of data according to the method proposed by Meinshausen and Bühlmann (2010). Probability of selection for each predictor is calculated over repeated variable selections.

An estimate of MB can be generated analogously by building an ensemble of regression optimal sets ℰ^opt^ identified without considering generalisability (i.e, let ℰ^gen^ = {1, …, *P*} ^2^ in the above procedure). The differences in the importance scores calculated based on ℰ^genopt^ and ℰ^opt^ highlights the predictors that are in NSB.

While the algorithm of Pfister et al. (2021) is statistically principled, it suffers from large computational drawbacks. Generalisable sets are selected based on enumerating all possible subsets of predictors. This can incur formidable computational cost even when the total number *P* of predictors is moderately large. To handle such cases, the authors suggested either to pre-screen the predictors or to consider only some random subsets of certain sizes, thus effectively reducing the number of predictor sets being considered. However, determining appropriate numbers and sizes of predictor sets to examine requires prior knowledge, as well as a trade-off between computational efficiency and accuracy. The selections resulting from random subsets are also highly variable – this was addressed by incorporating stability selection that repeats the SR procedure and considers the averaged selections over several repetition (Meinshausen and Bühlmann, 2010) . However, stability selection is also computationally intensive. These limitations hinder the wide applicability of SR in biological data.

#### 7.1.3 StableMate

We propose a highly flexible computational framework, StableMate, to improve the original SR procedure. Instead of selecting SB by exhaustive testing predictor sets on the regression optimal and generalisable conditions, we first use a greedy approach, namely an improved stochastic stepwise variable selection (ST2*), to select the *K* most predictive predictor sets 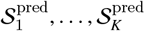 in MB (Xin and Zhu, 2012). Then, we narrow down our search by identifying the stable *and* predictive variables within 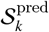 to obtain 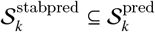, also using ST2*. This step is motivated by the fact that SB must be a subset of MB. This greedy approach avoids enumerating each subset of predictors. Selecting stable predictors in SB within the predictor sets MB also greatly reduces the search space and thus improvse computational efficiency. Similar ideas of search space reduction for finding stable predictors have been investigated in the domain of causal adaptation, e.g. Javidian et al. (2021); Rojas-Carulla et al. (2018).

##### Summary of methodological contributions

We emphasize the major modifications that distinguish StableMate from SR:

1. **Variable selection method:**
  - SR: selects stable and predictive predictor subsets by performing statistical tests on the regression optimal and generalisable conditions.
  - StableMate: employs ST2* to search for a predictor set that maximize an objective function quantifying stability or prediction ability.
  - Improvement: avoids the ambiguity in deciding appropriate significant thresholds for statistical tests, as well as the need of randomly sampling predictors subsets to test.
2. **Ensemble building:**
  - SR: randomly samples predictor subsets to test, and constructs an ensemble by the subsets that pass the tests.
  - StableMate: uses ST2* as a stochastic selector, and builds an ensemble by repeating the ST2* process.
  - Improvement: follows as a consequence of changing variable selection method.
3. **Order of selection**:
  - SR: selects stable predictor subsets first and then tests selected subsets for their prediction ability.
  - StableMate: runs ST2* to select sets of most predictive variables first, and then within each predictive set, runs ST2* to select a stable and predictive set.
  - Improvement: Stability is more difficult to assess compared to prediction ability since the former involves an additional environment factor to consider. Therefore, StableMate improves computational efficiency by reducing the search space of stable and predictive sets down from all predictors to predictive sets only. This modification is justified by the fact that a stable *and* predictive set must be a subset of a predictive set.

Other than the differences listed above, we also contributed in developing ST2* based on ST2 originally proposed by (Xin and Zhu, 2012). We not only improved the accuracy of ST2 (Supplementary Figure S2), but also introduced the concept of *pseudo-predictor* to create an automatic benchmark for the selection of true predictors in ensemble learning (see the paragraph below and also Supplementary Methods 7.1.4 for details).

##### Generating stable and predictive ensembles

To select the sets 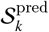and 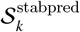, we developed a new version of stochastic stepwise variable selection ST2 from Xin and Zhu (2012), which we refer to as ST2*. Similar to stepwise selection, ST2* runs a greedy algorithm to search the subset of predictors maximising or minimising an objective function. Stochastic stepwise variable selection is designed to actively avoid suboptimal solutions, hence addressing a known limitation of stepwise selection. We postpone the description of our ST2* algorithm to Supplementary Methods 7.1.4 to first focus on how we use ST2* to generate ensembles of 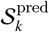 and 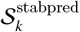 sets.

First, to select predictive sets 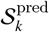, *k* = 1, …, *K*, we aim at maximising the objective function denoted obj^pred^, defined as the Negative Bayesian Information Criterion (NBIC) (see equation (10) in Supplementary Methods 7.1.5). For the purpose of statistical inference, we propose to include a pseudo-predictor indexed by *P* + 1. This pseudo-predictor is purely nominal: its inclusion in a regression model does not change the model fitting nor influences the objective function (see details in Section 7.1.5). We run ST2* *K* times to maximise obj^pred^. Each iteration results in a predictor set 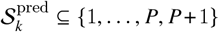. Note that 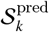 may include the pseudo-predictor, which will be useful for statistical inference. Since ST2* is stochastic, each run will result in a potentially different predictor set. We denote 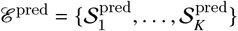 the resulted ensemble of *K* predictive predictor sets.

Next, for each *k* iteration we use ST2* to select stable predictors within 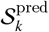 by maximising the objective function obj^stab^ defined as the Negative Prediction Sum of Squares (NPSS) (see equation (11) in Supplementary Methods 7.1.5). This enables us to identify an ensemble of stable and predictive sets 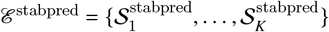, where each 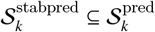.

##### Importance scores

After obtaining the ensembles ℰ^pred^ and ℰ^stabpred^, we evaluate the importance of each individual predictor. For *p* = 1, …, *P* + 1, we define the importance score 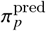 measuring the predictive ability of the predictor *p* (without considering its stability yet) as

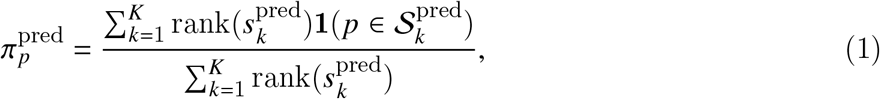

where 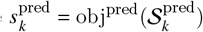 is the NBIC obtained on the predictive set 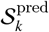, and 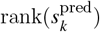 is the rank, in ascending order, of 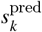 among 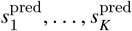 (ties are assigned a rank equals to the average of neighbouring integer ranks). The importance score 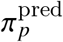 measures how frequent a predictor *p* is included in the 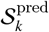 sets (large weights represent highly predictive sets).

To measure the stability of each predictor *p* within 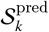, we then calculate the importance score 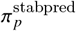 by considering both NBIC and NPSS, that is,

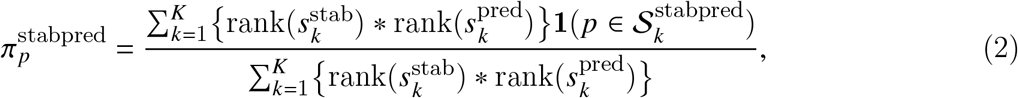

where 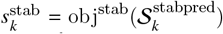 is the NPSS obtained on the stable and predictive set 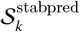, and 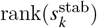 is the rank, in ascending order, of 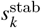 among 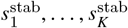.

As we consider the product of 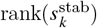 and 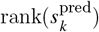 to weight the 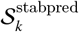 sets, a predictor *p* with a high score 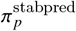 requires to be included more frequently in the 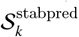 sets with both high prediction score *s*^stab^ and high stability score *s*^pred^. Therefore, a direct measurement of stability 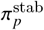 is obtained by adjusting 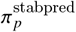 by normalizing according to the predictive ability of the predictor. We propose a measurement that is an analogy to conditional probability:

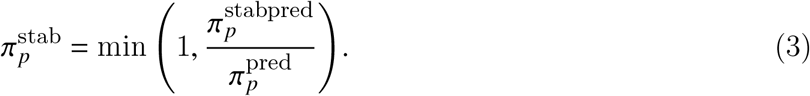

Intuitively, we measured the likely stability of predictor *p* when *p* is predictive. This score is output in our variable selection plot.

##### Identification of the blankets MB, SB and NSB

We use a bootstrapping approach to identify MB, SB and NSB by comparing the bootstrap distributions of the importance scores 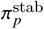 and 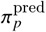 for all predictors *p* = 1, …, *P*, to the bootstrap distributions of the importance scores 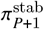 and 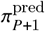 of the pseudo-predictor. Predictors with 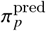 significantly larger than 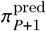 are considered predictive and are hence included in MB. Predictors in MB with 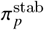 significantly larger than 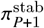 are considered as stable and predictive, and included in SB. Predictors in the MB with 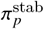 significantly smaller than 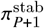 are considered as unstable but predictive, and are included in NSB. The remaining predictors are either considered outside the MB, or within MB but their stability cannot be ascertained by data at hand.

More specifically, for *b* = 1, …, *B*, where *B* is the total number of bootstrap iterations, we sample with replacement, *K* indices from {1, …, *K*} and denote the resample as 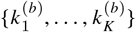. Let 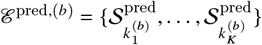 and 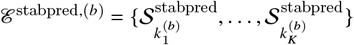 denote the *b*th bootstrap resample of ℰ^pred^ and ℰ^stabpred^, respectively. Then, for the predictors *p* = 1, …, *P*, we define the significance of stability based on the probability that 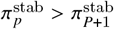 as

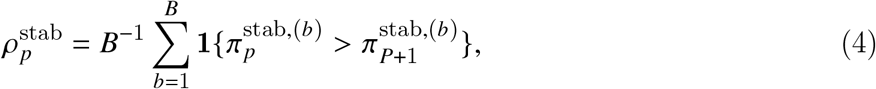

where the 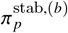 are the bootstrap versions of the importance scores 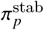 in (3). Similarly, we define the significance of unstable 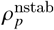 and of predictive ability 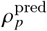 of the predictor *p* as in (4), as the estimates of the probabilities of 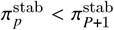 and 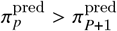, respectively.

Finally, a predictor *p* with 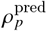 larger than a fixed significance threshold (e.g., 0.95) is selected as a predictor in MB. If such predictor *p* also has 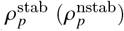 larger than this threshold, then it is selected as a predictor in SB (NSB respectively).

##### Final regression model

The identification of MB, SB and NSB is useful for interpreting the role of the different predictors in the regression model. It is then natural to use the predictors in SB to build a regression model that is generalisable to unseen environments. We use an ensemble approach for this purpose. Let 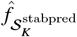 denote the regression model fitted by regressing *Y* on the predictor set 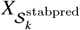 . Then, our aggregated model is defined as the weighted average of the 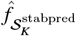, that is,

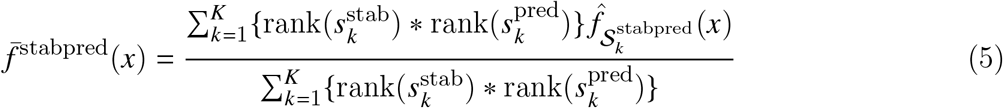

where the NBIC 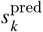 and the NPSS 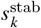 and their rank 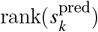 and 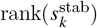 are defined as in equations (1) and (2).

Another version of aggregated model 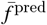 is the model trained without considering stability, defined by equation (5), but with 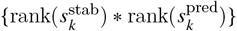 and 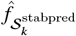 replaced by 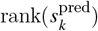 and 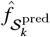, respectively. We compared the prediction performance of 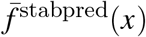 and 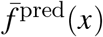 in Case study 2 (Section 2.3).

StableMate pseudo-code is presented in Algorithm 1.

###### Algorithm 1

StableMate

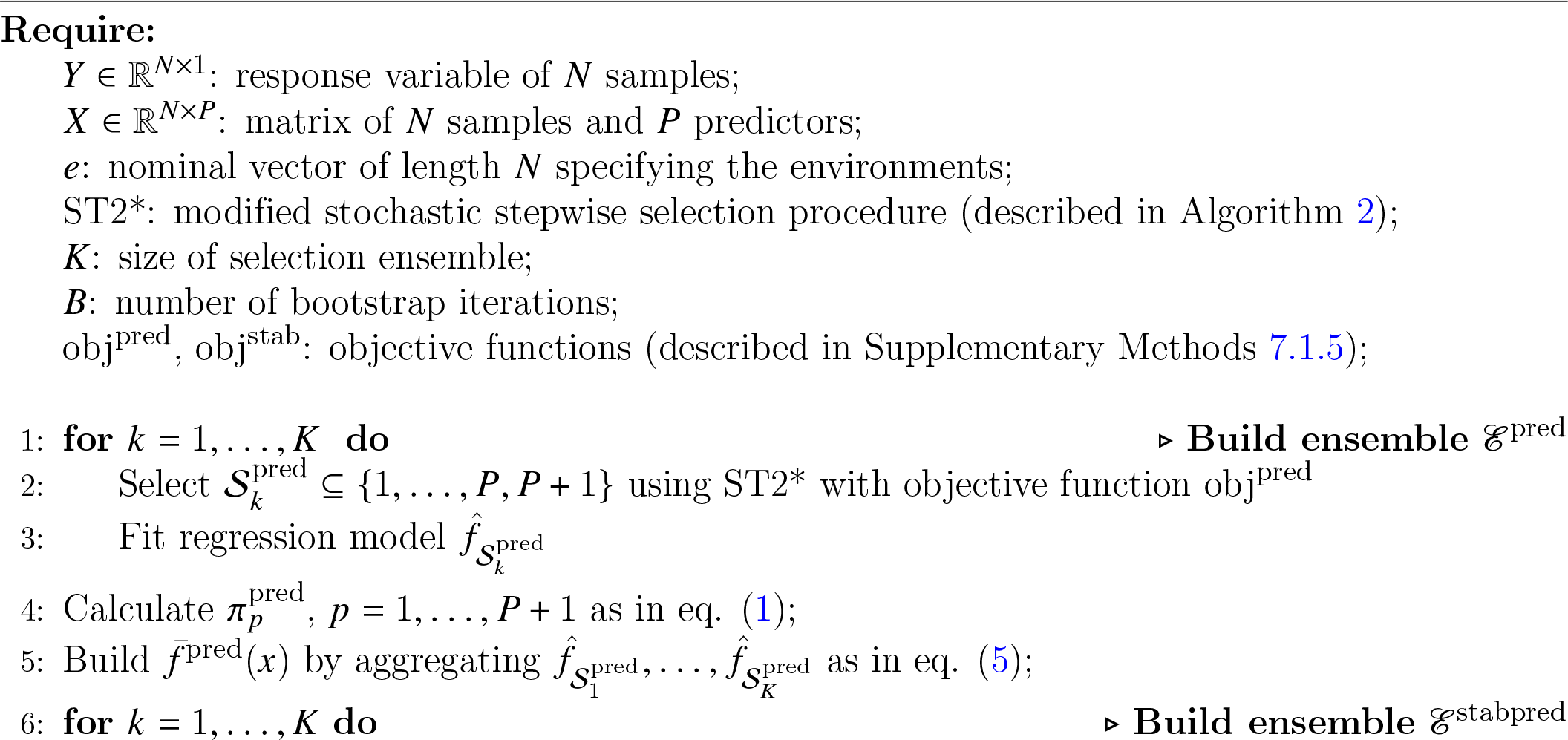
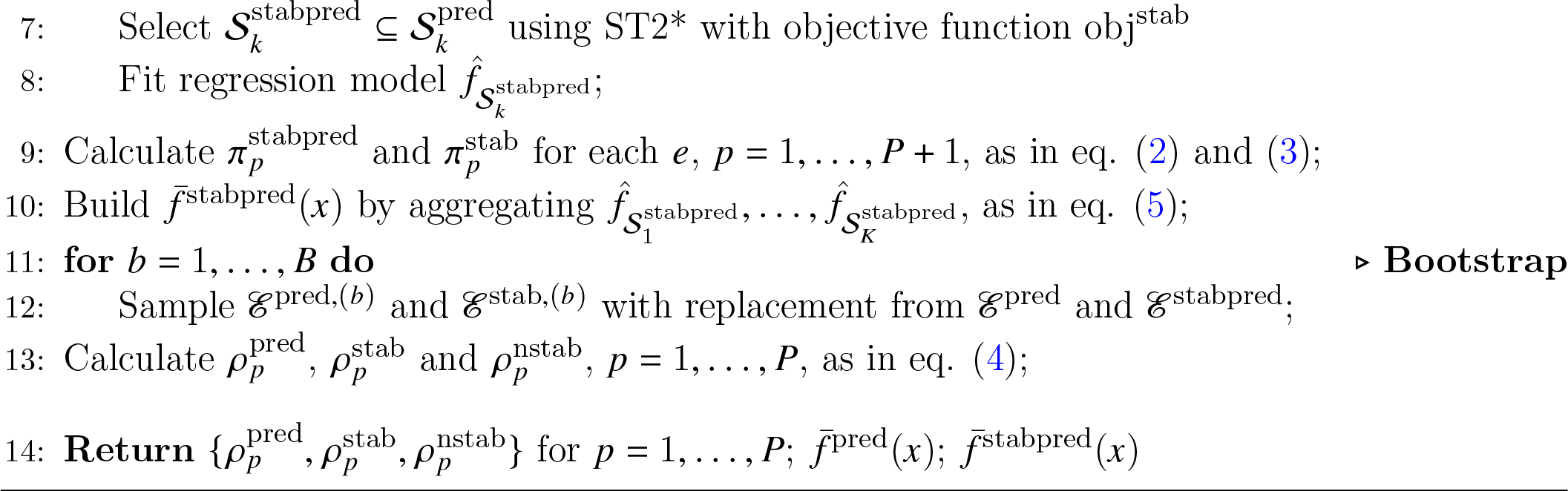

#### 7.1.4 ST2*: a new stochastic stepwise variable selection procedure

Our proposed ST2* plays an important part to select stable and predictive predictor sets, as we describe in this section.

ST2* improves Stochastic Stepwise Variable Selection (ST2, Xin and Zhu (2012)) to fit into the SR framework. Recall that a classical stepwise variable selection looks for predictor sets that maximise some objective function obj, e.g., the Bayesian Information Criterion (BIC), in a greedy search. It combines forward and backward steps iteratively. In a forward (respectively, backward) step, we add (respectively, delete) one predictor at a time to the current model that best increase the value of the objective function. The stepwise variable selection is complete when the objective function is not longer improved.

While stepwise variable selection is scalable to high dimensionality, it fails to select important predictors when they are highly correlated (Smith, 2018). This is a strong limitation in our case as we expect some underlying causal and correlated structures of our predictors. ST2 was proposed by randomly select a number of candidate predictor sets and then choose the best set to add to or remove from the current model for each forward or backward step (Xin and Zhu, 2012). We refer to candidate predictor set as a *proposal* and the number of predictors in a proposal as the *proposal size*. For each forward / backward step, ST2 generates proposals of the same size. However, our simulation studies showed that an ill-suited proposal size can cause ST2 to stop prematurely. We therefore proposed a new ST2* procedure that randomises proposal sizes.

##### Proposal size sampling in forward selection in ST2*

Let 𝒮 ^*pool*^ be the predictor pool wherein an optimal predictor set is searched, and 𝒮 ^∗^ the set of predictors in the current model. At a forward step, we generate a proposal 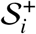 by sampling without replacement 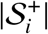| predictors from 𝒮 ^*c*^ = 𝒮 ^*pool*^ − 𝒮 ^∗^, the set of predictors not in the current model, for *i* = 1, …, *η*^+^, where *η*^+^ denotes the total number of proposals to generate at the current step (we will discuss shortly how to select *η*^+^).

The proposal size 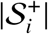 is sampled from a shifted beta-binomial distribution

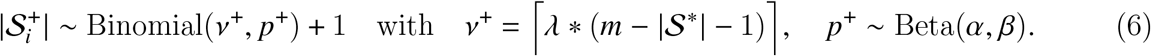

where *v*^+^ is the proposal size, *p*^+^ is the success probability of the binomial distribution, and *λ, m, β* and *α* are hyperparameters:

- *m* is a positive integer that satisfies 0 < *m* ≤ | 𝒮 ^*pol*^ | and represents the maximum number of predictors allowed in a model. If | 𝒮 ^∗^| ≥ *m*, then we skip the current forward step to the next backward step.
- *λ* is a fine tuning parameter with 0 < *λ* ≤ 1. We define *v*^+^ so that 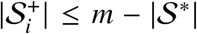 – this upper bound represents the maximum number of predictors that can be added when *λ* = 1.
- *α* and *β* control the shape and the scale of the beta distribution.

By default, we set *λ* = 0.5, *α* = 1 and *β* = 5 to allow for left skewed sampling of proposal sizes. Our rationale is to propose small steps for local search, while allowing large steps with relatively smaller probabilities to reduce the likelihood of local optimal. The choice of *m* is data specific. We set *m* = | 𝒮^*pol*^ |, which seemed to work well in our simulation and case studies.

Among the forward proposals 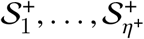, we add the proposal to the current model that best improve the objective function obj to complete the forward step.

##### Proposal size sampling in backward selection in ST2*

The backward step is described analogously to the forward step. Briefly, we generate *η*^−^ backward proposals 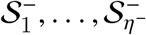 by sampling from 𝒮^*c*^. The proposal size 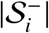 is sampled by

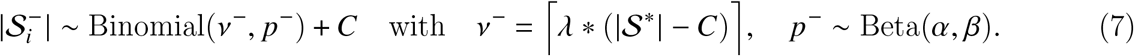

where *C* = max(|𝒮^∗^| − *m*, 1) is the number of predictors that must be removed from 𝒮^∗^ as restricted by . If the model is empty and there are no more predictors to remove, we directly enter the next forward step. Both backward and forward steps shares the same hyperparameters in eq. (6). Since we cannot remove more predictors than those in the current model 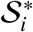, our definition of *v*^−^ restricts that 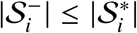 when *λ* = 1.

##### Number of proposals to sample in forward selection

To complete our ST2* procedure, we need to determine how the total number of proposals *η*^+^ (forward step) and *η*^−^ (backward step) are selected.

The original ST2 performs a grid search to select a best *η*^+^, which is computationally intensive. A small change in the tuning of *η*^+^ originally proposed by Xin and Zhu (2012) results in a large change in *η*^+^. However, we found that in practice, ST2 was not very sensitive to the *η*^+^ value, as long as it was not extremely large or small. This motivated our following heuristic approach for selecting *η*^+^.

ST2* chooses the number *η*^+^ of proposals at each forward step as

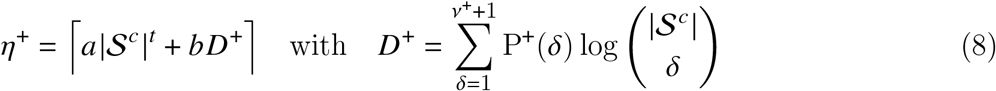

where P^+^(*δ*) is the probability of generating a proposal of size *δ* from 𝒮^*c*^ (predictors not in the current model), and *v*^+^ + 1 is the maximum size of the proposals that can be generated. P^+^(*δ*) and *v*^+^ are calculated as in eq. (6). The binomial coefficient 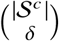 represents the amount of unique proposals of size *δ* that can be potentially generated. *D*^+^ is defined as a weighted average of the logarithm of binomial coefficients. If proposals of size *δ* are more likely to be generated, then its abundance influences more on *D*^+^ and hence *η*^+^. The more unique proposals are available, the larger *η*^+^ is. In the extreme case where P^+^(*δ* = |𝒮^*c*^ |) = 1, *η*^+^ = 1 is all we need for sampling 𝒮^*c*^, which corresponds to the only proposal available. If P^+^(*δ* = 1) = 1, then each predictors in |𝒮^*c*^ | can be generated as a unique proposal by itself. In this case, it would be better to make more than one proposal. *a* |**S**^−^| ^*t*^ defines a lower bound of *η*^+^ that can be flexibly tuned in case *D*^+^ becomes too small due to the logarithm. *a, b* and *t* are new hyperparameters. We found in our simulation studies that ST2* procedure worked reasonably well with *a* = 1, *b* = 1 and *t* = 0.5.

##### Number of proposals to sample in a backward selection

Similarly to the forward selection, we define *η*^−^ as follows,

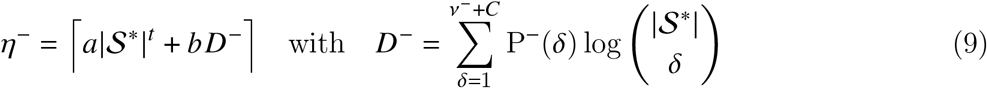

where P^−^(*δ*) is the probability of generating a proposal of size *δ* from 𝒮^∗^ (predictors in the current model), *v*^−^ + *C* is the maximum size of proposals that can be generated. We refer the calculation of P^−^(*δ*) and *v*^−^ + *C* to eq. (7). We use the same parameters *a, b, t* as in the forward selection.

We present the pseudocode for our ST2* procedure in Algorithm 2. Since some preliminary filtering of predictors can be applied prior to ST2* – for example by using random Lasso as we propose, we denote 𝒮^pool^ the pool of filtered predictors. Recall that ST2* aims to select 𝒮^∗^ ⊆ 𝒮pool to maximise some objective function obj. In StableMate (Algorithm 1), to build ensemble ℰ^pred^ with ST2*, we set by default 𝒮^pool^ = {1, …, *P* +1} when no predictor filtering is needed (see Supplementary Section 7.1.6 otherwise); to build ensemble ℰ^stabpred^ with ST2*, we have 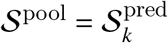, since each 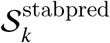 is restricted to be a subset of 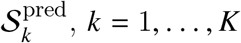.

###### Algorithm 2

ST2* stochastic stepwise selection

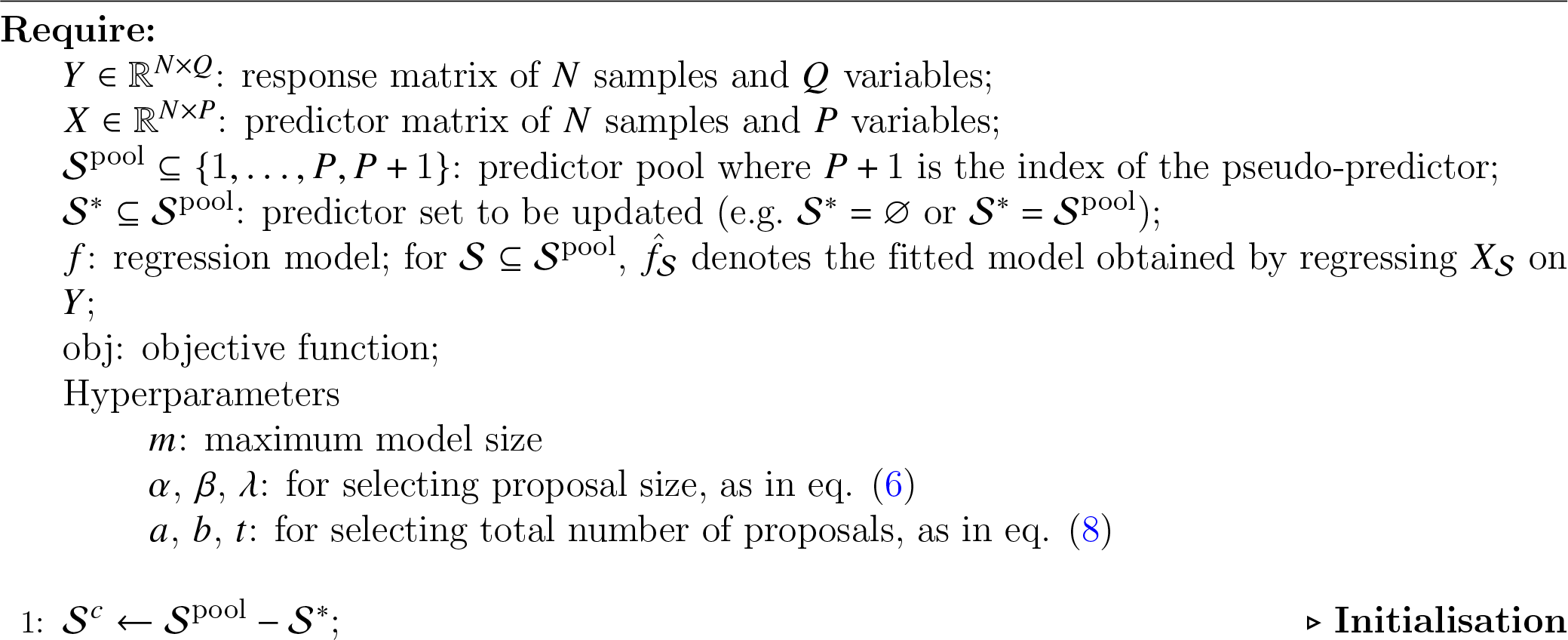
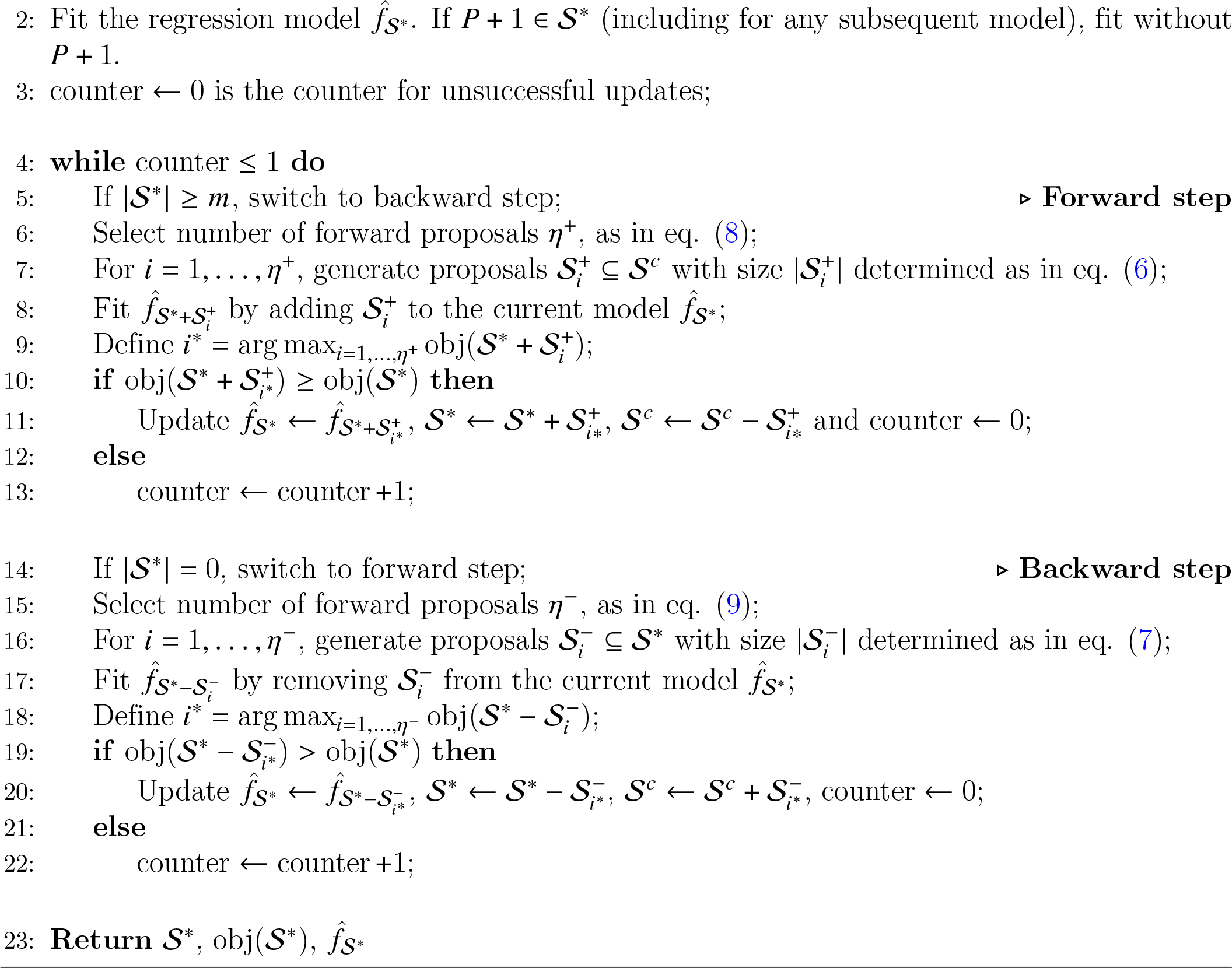

#### 7.1.5 Objective functions in StableMate and definition of the pseudo-predictor

We define the objective functions in StableMate when using Ordinary Least Square regression as the default regressor of ST2* for selecting stable and predictive predictor sets.

Denote 𝒮 ⊆ {1, …, *P* + 1} a predictor set where *P* + 1 is the index of the pseudo-predictor. For environment *e* ∈ ℰ, *N*^*e*^ denote the number of samples observed in *e* and *N* = Σ^*e*∈ℰ^ *N*^*e*^ denote the sample size. Denote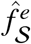 the regression model fitted using data in environment *e* and using 𝒮 as predictors. Denote 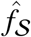 the regression model fitted using data pooled across all environments and using 𝒮 as predictors.

The pseudo-predictor *P* + 1 is an index that can be added or removed from 𝒮. When the pseudo-predictor is included in 𝒮, the regression models are fitted on all predictors in 𝒮 but without the pseudo-predictor. However, we treat the pseudo-predictor as if it was used to fit the model, similar to the other predictors in 𝒮. Therefore, the pseudo-predictor can be selected by ST2* but does not influence the model fitting nor the objective function, which depends on 𝒮 through the fitted model.

To measure the predictive ability of the predictor set 𝒮, we define the Negative Bayesian Information Criterion (NBIC) obj^pred^(𝒮) for 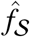 as

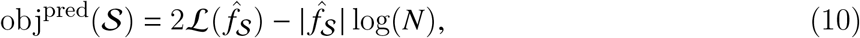

where 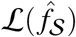 denotes the log-likelihood of 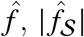 the model complexity and *N* is the pooled sample size. obj^pred^ attains the largest value when 𝒮 is the Markov blanket of the response, hence this criterion can be used to identify MB (see Supplementary Methods 7.1.1).

To measure the stability of the predictor set 𝒮, we define the objective function obj^stab^(𝒮) as the Negative Prediction Sum of Squares (NPSS) of all 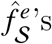 as

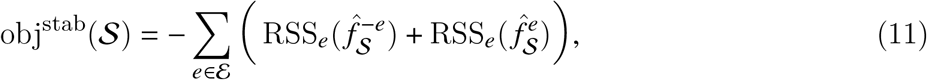

where 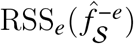 denotes the residual sum of squares from the regression function 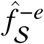 trained without the environment *e* to data observed in the environment *e*. A small 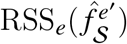 implies that 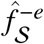 is more generalisable to the new environment *e*. Hence if the predictor set 𝒮 is truly stable so that all 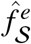 with *e* ∈ ℰ tend to be generalisable to other environments, then RSS^*e*^values will be small and hence obj^stab^(𝒮) large. Since the within environment measure of predictive ability 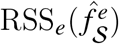 always decreases with increasing model complexity, we add this term in eq. 11 to ensure that 𝒮^stabpred^ ⊆ 𝒮^pred^ obtains the highest obj^stab^(𝒮^stabpred^) at 𝒮^stabpred^ = 𝒮^pred^ if there is no environmental effect that perturbs the prediction.

We have extended these objective functions in StableMate for Generalized Linear Model by replacing RSS in (11) by the sum of deviance.

#### 7.1.6 Pseudo-predictors in Lasso pre-screening in StableMate

Since pseudo-predictors are indexes with no real form (as described in Supplementary Methods 7.1.5), Lasso selection is not applicable to them. Therefore, we included a pseudo-predictor in all pre-filtered sets to calculate importance scores of selections as in equation (1). To benchmark selections as in equation (4), the importance score of the pseudo-predictor needs to adjust for the fact that the true predictors are pre-filtered but not the pseudo-predictor. Pseudo-predictors are defined as predictors at the boundary of selections where they can be either selected or not. If pseudo-predictors are selected by Lasso, they are expected to be selected with confidence similar to the *p*th important predictor of Lasso, which is also at the boundary. Therefore, we propose to downscale the importance score of a pseudo-predictor by multiplying it with the *p*^*th*^ largest selection frequency of Lasso to mimic Lasso selection on that pseudo-predictor.

### 7.2 Simulation studies

We conducted simulation studies for the following purposes:

- To generate a toy example to illustrate the StableMate analysis in Section 2.1;
- To compare the performances of 3 methods in selecting highly predictive variables, including the new ST2* algorithm used in StableMate, the ST2 algorithm in Xin and Zhu (2012) and the original SR algorithm in Pfister et al. (2021). The results are reported in Supplementary Methods Section 7.2.2.
- To compare the performances of StableMate and the original SR algorithm in selecting stable and environment-specific predictors. The results are reported in Supplementary Methods Section 7.2.3.
- To compare the performances of StableMate, SR, ordinary least squares regression, Lasso regression (OLS, Tibshirani 1996) and random forest (RF, Breiman 2001) in building regression models generalisable to unseen environments. The results are reported in Supplementary Methods Section 7.2.3.

#### 7.2.1 Model and methods

As in Pfister et al. (2021), we simulated data from structural causal models to benchmark our method. To construct an structural causal model, we first need to construct a directed acyclic graph (DAG), which is a graph representation of the underlying causal structure of all variables. Specifically in our setting, a node on DAG represents either a variable (a response and predictors) or a source of environmental perturbation, whereas a directed edge between two nodes represents the causal relations between them. Then, we simulate observations on each variable according to the DAG structure.

To simulate a DAG with environmental interventions, we used a two-step procedure as follows. Firstly, we use the *rgraph* function from the R package `gmat` to randomly generate a DAG. For our regression analysis, we randomly choose a node (variable) as the response *Y* ^*e*^, with the remaining variables as predictors 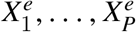 . Then, to simulate environmental interventions, we add to the DAG a set of exogenous nodes *ξ*^1^, …, *ξ*^*E*^ (the number *E* is proportional to the number nodes in DAG; see Section 7.2.2), each randomly assigned as a parent of one of the predictor nodes. These exogenous nodes simulate the environmental interventions exogenously influencing the causal structure of the predictors. Given a DAG, we can then identify three sets of predictors, namely the Markov blanket (MB), i.e. the predictive variables, the stable blanket (SB), i.e. the stable predictors and the non-stable blanket (NSB), i.e. the environment-specific predictors. MB and SB have been defined in Supplementary Methods 7.1.1 and NSB is the set difference between MB and SB. For the relationship between the MB, SB and NSB and how to identify them on a given DAG, see Pfister et al. (2021).

Given a DAG specifying the causal relations of variables and the environmental interventions, as in Pfister et al. (2021), we simulate observations from 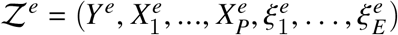 using the following linear model

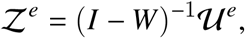

where *I* is a *P* by *P* identity matrix, *W* is a weight matrix representing the causal structure specified by DAG, and 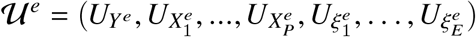 is a multivariate normal random vector. Let 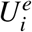 denote a generic element of 𝒰^*e*^ . We assume that

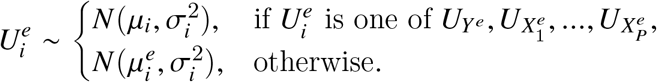

That is, the endogenous 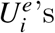 are identically distributed across environments but the exogenous ones have environment-specific distributions.

To simulate samples from environments *e* ∈ ℰ, we first generate the weight matrix *W* = (*W*^*i, j*^) and the distributional parameters 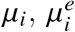 and 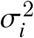 as follows. To simplify the notation, let *Z*^*i*^ and *Z* ^*j*^ denote two generic nodes. Then *W*^*i, j*^ is defined by

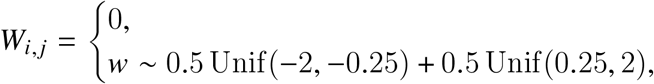

Then, we generate

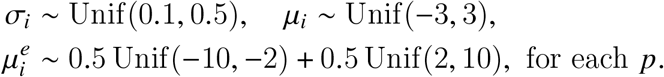

Once these parameters *W, μ*^*i*^, 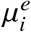 and 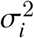 are generated, we treat them as deterministic and simulate from 𝒰 ^*e*^ for all *e* ∈ ℰ.

We keep only copies of *Y* ^*e*^ and 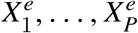 as simulated data. The simulated environmental interventions are discarded as they are not observable in practice.

#### 7.2.2 Benchmark study for predictivity selection

In this section, we benchmark the performance of our ST2* algorithm in selecting predictive variables, i.e. predictors in the underlying MB, in a single environment (i.e. no stability selection is involved), against the ST2 algorithm in Xin and Zhu (2012) and the original SR algorithm in Pfister et al. (2021).

We considered 3 DAGs with increasing number of nodes. In addition to 1 response, the first DAG has 50 predictors, 30 environmental interventions and 10 percent of all possible edges connected; the second DAG has 100 predictors, 60 interventions and 5 percent of all possible edges connected; the third DAG has 200 predictors, 120 interventions and 2.5 percent of all possible edges connected. From each DAG, we simulated 100 single-environment datasets each with 300 samples.

Both ST2 and ST2^∗^ are ensemble-based approaches and a key tuning parameter is the number of selections (predictor sets) per ensemble. In our simulation study, we included 100 selections in each ensemble. As for other tuning parameters, ST2^∗^ was run with the default parameters as in Section 7.1.4, while ST2 was tuned so that its running time was approximately equal to the running time of ST2^∗^. This is the computation of ST2 will be much more time-consuming if we use their default parameters. To quantify the performance of SR, we used the stability selection strategy as in Meinshausen and Bühlmann (2010) (see 7.1.2). That is, for each simulated dataset, we repeated the SR procedure for 100 times. In each run, to reduce the high computational cost of SR, we did a Lasso pre-screening to reduce the number of variables. In particular, for a dataset with *P* predictors, we pre-selected *P*/5 predictors as input to the SR algorithm. Then, the input predictors were randomly subsampled 20*P* times to generate 20*P* predictor sets. The negative Bayesian information criterion (NBIC) defined in Supplementary Methods 7.1.5 was used to assess the prediction ability of each predictor set. Predictors within the sets that were founded significantly predictive (with significance level 0.01) were viewed as an estimate of MB. This SR procedure of MB selection was then repeated 100 times and a selection probability of each predictor was calculated.

To evaluate how accurately each method selected predictors in MB, we computed the area under the curve (AUC) as follows. For ST2^∗^ and ST2, AUC were calculated based on the prediction score defined by (1) in Section 7.1.3. For SR, the AUC was computed based on the selection probabilities returned by the stability selection procedure as described above. Since ST2 and ST2^∗^ had similar performance measured by their AUC, we further compared them by the mean NBIC. The results are summarised in Supplementary Figure S2, showing that ST2* was the most accurate method in selecting MB among all three methods and it had higher efficiency to achieve the optimal NBIC compared to ST2.

#### 7.2.3 Benchmark study for stability selection and model generalisability

We first benchmark the performance of StableMate in selecting stable and environment-specific predictors, against the original SR algorithm in Pfister et al. (2021). Then we benchmark the performance of the regression model built by StableMate using stable predictors, against the original SR algorithm, OLS regression, Lasso regression (Tibshirani, 1996) and RF (Breiman, 2001). From each of the 3 DAGs described in Section 7.2.2, we simulated 100 datasets, each containing 1200 samples from 4 environments (300 samples per environment).

To implement StableMate, we used the default parameters as described in Section 7.1.3. SR was implemented with Lasso pre-filtering as described in Supplementary Methods 7.2.2. After building an ensemble of predictor sets with significantly high prediction ability, SR conducted a stability test based on the Chow-test (Chow, 1960) with 0.01 significance level to build an ensemble of stable predictor sets. Predictors within the stable predictor sets were selected as in SB. Predictors that are in MB but not in SB were selected as in NSB. Again, we repeat this SB (or NSB) selection procedure for 100 times, and a selection probability of each predictor was calculated.

As in Section 7.2.2, we benchmarked StableMate and SR selections for SB using AUC, this time based on the stability scores (2) in Section 7.1.3 and selection probabilities of SB predictors. Since StableMate does not generate selection importance scores for NSB predictors to calculate AUC for selections, we compared instead the balanced accuracy of NSB selections (defined as the average of sensitivity and specificity) between StableMate and SR.

Finally, to compare the prediction performance of StableMate to SR, OLS, Lasso and random forest, we trained each method in the first three environments and computed the negative mean squared errors of the fitted regression model applied to the fourth environment. We implemented random forest using the R package `randomForest` with the default tuning parameters. The Lasso regression was tuned with cross-validation. The results are summarised in Supplementary Figure S1. From there we can see that StableMate and SR achieved superior prediction in unseen test environments compared to the other regression methods. Furthermore, StableMate outperformed SR, achieving higher prediction and variable selection accuracy with reduced computational cost.

